# Mitochondrial Apolipoprotein MIC26 is a metabolic rheostat regulating central cellular fuel pathways

**DOI:** 10.1101/2023.12.01.569567

**Authors:** Melissa Lubeck, Ritam Naha, Yulia Schaumkessel, Philipp Westhoff, Anja Stefanski, Patrick Petzsch, Kai Stühler, Karl Köhrer, Andreas P. M. Weber, Ruchika Anand, Andreas S. Reichert, Arun Kumar Kondadi

## Abstract

Mitochondria play central roles in metabolism and metabolic disorders such as type 2 diabetes. MIC26, a MICOS complex subunit, was linked to diabetes and modulation of lipid metabolism. Yet, the functional role of MIC26 in regulating metabolism under hyperglycemia is not understood. We employed a multi-omics approach combined with functional assays using WT and *MIC26* KO cells cultured in normoglycemia or hyperglycemia, mimicking altered nutrient availability. We show that MIC26 has an inhibitory role in glycolysis and cholesterol/lipid metabolism under normoglycemic conditions. Under hyperglycemia, this inhibitory role is reversed demonstrating that MIC26 is critical for metabolic adaptations. This is partially mediated by alterations of mitochondrial metabolite transporters. Furthermore, *MIC26* deletion led to a major metabolic rewiring of glutamine utilization as well as oxidative phosphorylation. We propose that MIC26 acts as a metabolic ‘rheostat’, that modulates mitochondrial metabolite exchange via regulating mitochondrial cristae, allowing cells to cope with nutrient overload.

---

Mitochondria, Apolipoproteins, MIC26, MICOS complex and Fatty acid metabolism

## Introduction

The increasing prevalence of global obesity is a huge biological risk factor for development of a range of chronic diseases including cardiovascular diseases, musculoskeletal and metabolic disorders (Collaborators *et al*, 2017). At the cellular level, obesity is associated with DNA damage, inflammation, oxidative stress, lipid accumulation and mitochondrial dysfunction (Włodarczyk & Nowicka, 2019). Mitochondria play central roles in anabolic and catabolic pathways (Spinelli & Haigis, 2018) and as a consequence mitochondrial dysfunction is associated with a variety of metabolic diseases such as type 2 diabetes mellitus (T2DM) (Szendroedi *et al*, 2011). Mitochondrial dysfunction is often linked to abnormal mitochondrial ultrastructure (Eramo *et al*, 2020; Kondadi *et al*, 2020b; Zick *et al*, 2009) and abnormal mitochondrial ultrastructure was also associated with diabetes (Bugger *et al*, 2008; Xiang *et al*, 2020). Mitochondria harbour two membranes, the mitochondrial outer membrane (OM) and the inner membrane (IM). The part of the IM closely apposed to the OM is termed the inner boundary membrane (IBM) whereas the IM which invaginates towards the mitochondrial matrix is termed the cristae membrane (CM). Crista junctions (CJs) are pore-like structures around 12 to 25 nm in diameter, separating the IBM and CM, and are proposed to act as diffusion barriers for proteins and metabolites (Frey & Mannella, 2000; Mannella *et al*, 2013). Formation of CJs depends on Mic60 (Fcj1, Mitofilin, IMMT) which was shown to be located at CJs regulating cristae formation in concert with the F_1_F_O_ ATP synthase (Rabl et al. 2009). Mic60 is a subunit of the ‘mitochondrial contact site and cristae organising system’ (MICOS) complex (Harner *et al*, 2011; Hoppins *et al*, 2011; von der Malsburg *et al*, 2011) which consists of seven proteins organised into two subcomplexes: MIC60/MIC19/MIC25 and MIC10/MIC26/MIC27 with MIC13 stabilizing the MIC10 subcomplex in mammals (Anand *et al*, 2016; Guarani *et al*, 2015; Urbach *et al*, 2021). MIC26/APOO harbours an apolipoprotein A1/A4/E family domain and therefore was classified as an apolipoprotein (Koob *et al*, 2015; Lamant *et al*, 2006). Traditionally, apolipoproteins mediate lipid and cholesterol metabolism by facilitating the formation of lipoproteins and regulating their distribution to different tissues via the blood stream (Mehta & Shapiro, 2022). Initially, MIC26 was identified as a protein of unknown function in cardiac transcriptome of dogs fed with high-fat diet (HFD) (Philip-Couderc *et al*, 2003) and was incorrectly assumed to exist as a 55 kDa O-linked glycosylated protein as it was immunopositive to a custom-generated MIC26 antibody in samples of human serum, heart tissue and HepG2 cell line (Lamant *et al*., 2006). However, the recombinant protein was only observed at the expected size of 22 kDa (Lamant *et al*., 2006) and it was later shown that this 22 kDa form is located to mitochondria (Koob *et al*., 2015; Ott *et al*, 2015). Moreover, using several cellular *MIC26* deletion models and different antibodies, we showed recently that MIC26 is exclusively present as a 22 kDa mitochondrial protein and not as a 55 kDa protein (Lubeck *et al*, 2023). In light of these findings, the primary physiological function of MIC26 in diabetes is linked to its role in the mitochondrial IM and not to an earlier proposed secreted form of MIC26.

Mutations in *MIC26* were reported to result in mitochondrial myopathy, lactic acidosis and cognition defects (Beninca *et al*, 2021) as well as a lethal progeria-like phenotype (Peifer-Weiß *et al*, 2023). Interestingly, there is an intricate connection between MIC26 and metabolic disorders. Patients with diabetes (Lamant *et al*., 2006) and dogs fed with a HFD for 9 weeks (Philip-Couderc *et al*., 2003) showed increased *Mic26* transcripts in the heart. Accordingly, adenovirus-mediated human *MIC26* overexpression in mice, administered through the tail vein, led to increased levels of triacylglycerides (TAG) in murine plasma, when fed with HFD, and TAG accumulation in the murine liver (Tian *et al*, 2017). In another study, MIC26 transgenic mice hearts displayed an increase of diacylglycerides (DAG) but not TAG (Turkieh *et al*, 2014) as in the previous described study (Tian *et al*., 2017) suggesting modulatory roles of MIC26 in lipid metabolism. Recently, in mitochondria-rich brown adipose tissue (BAT), downregulation of *Mic26* mRNA and protein levels were reported in diet-induced or leptin-deficient obese (ob/ob) murine models compared to the respective controls. Mice with an adipose tissue specific deletion of *Mic26* which were fed with a HFD gained more total body weight and adipose tissue fat mass than control mice (Guo *et al*, 2023). Hence, we hypothesize that MIC26 has an unidentified regulatory role under nutrient-enriched conditions. Therefore, in order to understand the function of MIC26, we used WT and *MIC26* KO cells as a model system under standard glucose culture conditions as well as excessive glucose culture conditions termed normoglycemia and hyperglycemia, respectively. We employed a multi-omics approach encompassing transcriptomics, proteomics and targeted metabolomics to investigate the pathways regulated by MIC26. We found that the function of MIC26 is critical in various pathways regulating fatty acid synthesis, oxidation, cholesterol metabolism and glycolysis. Interestingly, we observed an entirely antagonistic effect of cellular *de novo* lipogenesis in *MIC26* KO cells compared to WT cells depending on the applied nutrient conditions. This showed that the response to high glucose conditions is strongly dependent on the presence of MIC26. In addition, we found that cells deleted for *MIC26* displayed alterations of mitochondrial glutamine usage and oxidative phosphorylation. Overall, we propose that MIC26 is a unique mitochondrial apolipoprotein functioning as a mitochondrial fuel sensor that regulates central metabolic pathways to meet mitochondrial and thus cellular energy demands.

## Results

### Mitochondrial apolipoprotein MIC26 is selectively increased in cells exposed to hyperglycemia

There is a strong link between metabolic abnormalities and pattern of *MIC26* expression. Increased levels of *MIC26* transcripts were observed in diabetic patients (Lamant *et al*., 2006) and increased accumulation of lipids were found upon *Mic26* overexpression in the mouse (Tian *et al*., 2017; Turkieh *et al*., 2014). In order to understand the role of MIC26 in cellular metabolism, we used hepatocyte-derived HepG2 cells as the cellular model and generated *MIC26* KO cells using the CRISPR-Cas9 system. WT and *MIC26* KO cells were grown in standard (5.5 mM) and excessive concentrations of glucose (25 mM), defined as normoglycemic and hyperglycemic conditions respectively throughout the manuscript, for a prolonged period of three weeks to investigate long term effects of nutritional overload. Initially, we checked whether there is a difference in the amounts of various MICOS proteins in WT HepG2 cells grown in normoglycemia and hyperglycemia. Western blot (WB) analysis showed a significant increase of MIC26 and MIC27 along with MIC25 in cells grown in hyperglycemia compared to normoglycemia (**Fig 1A** **& B**). We did not observe any significant changes in the amounts of MIC19, MIC60, MIC10 and MIC13 in WT-Hyperglycemia (WT-H) compared to WT-Normoglycemia (WT-N) condition. This pointed to a specific role of the MICOS subunits, MIC26, MIC27 and MIC25 when cultured in hyperglycemia compared to normoglycemia. The significant increase of MIC27 and MIC25 observed in WT-H when compared to WT-N was abolished in *MIC26* KOs indicating a requirement of MIC26 in this response under nutrient-enriched conditions (**Fig 1A** **& B**).

**Figure 1.**
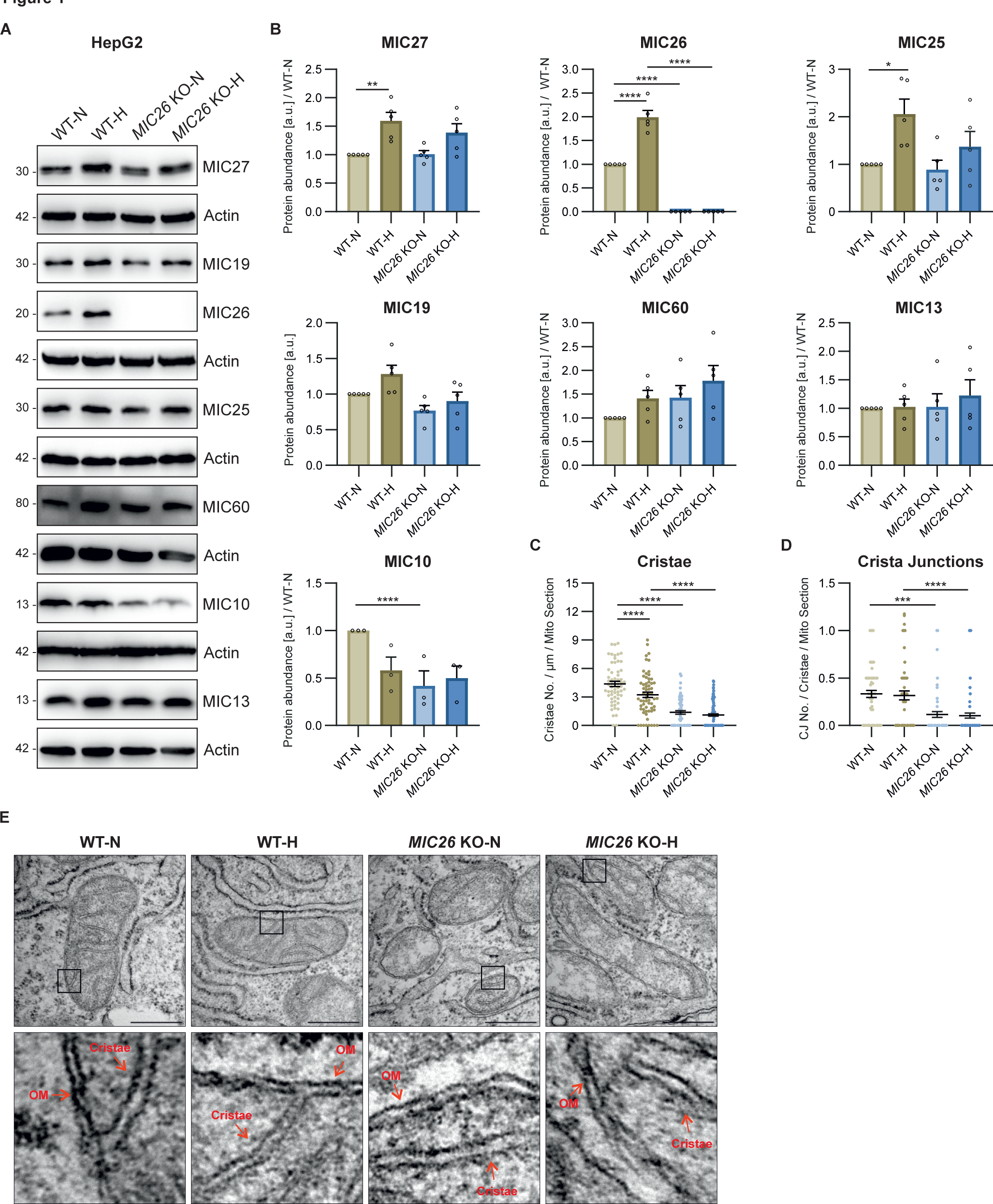
Mitochondrial apolipoprotein MIC26 is selectively increased in cells exposed to hyperglycemia. (A and B) Western blot analysis of all MICOS subunits from HepG2 WT and *MIC26* KO cells cultured in normo- and hyperglycemia (N = 3-5). Chronic hyperglycemia treatment leads to increased levels of MIC27, MIC26 and MIC25 in WT cells. Loss of MIC26 is accompanied by decreased MIC10 in normoglycemia. (C, D and E) Electron microscopy data including quantification of cristae number per unit length (µm) per mitochondrial section (C) as well as crista junctions per cristae per mitochondrial section (D), along with representative images (E) from HepG2 WT and *MIC26* KO cells cultured in normo- and hyperglycemia (N = 2). Loss of MIC26 led to decreased cristae number and crista junctions independent of normo- and hyperglycemia. Red arrows in lower row indicate outer membrane (OM) or cristae. Scale bar represents 500 nm. Data are represented as mean ± SEM (B, C and D). Statistical analysis was performed using one-way ANOVA with \**P* < 0.05, \*\**P* < 0.01, \*\*\**P* < 0.001, \*\*\*\**P* < 0.0001. N represents the number of biological replicates.

The MICOS proteins regulate the IM remodelling by working in unison to maintain CJs and contact sites between IM and OM (Anand *et al*, 2021). Still, deficiency of different MICOS proteins shows variable effects on the extent of CJs loss and cristae ultrastructure (Anand *et al*, 2020; Kondadi *et al*, 2020a; Stephan *et al*, 2020; Weber *et al*, 2013). MIC10 and MIC60 have been considered as core regulators of IM remodelling displaying severe loss of CJs (Kondadi *et al*., 2020a; Stephan *et al*., 2020). The extent of mitochondrial ultrastructural abnormalities upon *MIC26* deletion varies among different cell lines tested (Anand *et al*., 2020; Koob *et al*., 2015; Stephan *et al*., 2020). Therefore, we performed transmission electron microscopy (TEM) in WT and *MIC26* KO HepG2 cells which revealed a reduction of cristae content (cristae number per unit mitochondrial length per mitochondria) in *MIC26* KOs compared to WT cells in both nutrient conditions (**Fig 1C** **& E**). Thus, the loss of cristae was dependent on MIC26 and independent of the glucose concentration used in cell culture. In addition, there was a decrease in cristae number in WT cells grown in hyperglycemia compared to normoglycemia showing that higher glucose levels lead to decreased cristae density, which is a common phenotype in diabetic mice models (Bugger *et al*., 2008; Xiang *et al*., 2020). As the number of cristae were already decreased in certain conditions, we analysed the number of CJs normalised to cristae number and found that a significant decrease of CJs was observed in *MIC26* KOs independent of the nutrient conditions (**Fig 1D** **& E**). Overall, the loss of MIC26 leads to mitochondrial ultrastructural abnormalities accompanied by reduced number of cristae as well as CJs compared to WT cells (**Fig 1C-E**).

### Hyperglycemia confers antagonistic regulation of lipid and cholesterol pathways, in ***MIC26* KO vs WT cells, compared to normoglycemia**

In order to understand the role of MIC26 in an unbiased manner, we compared WT and *MIC26* KO cells cultured under respective nutrient conditions by employing quantitative transcriptomics and proteomics analyses. A total of 21,490 genes were obtained after the initial mapping of the RNA-Seq data, of which 2,933 were significantly altered in normoglycemic *MIC26* KO compared to WT cells (fold change of ±1.5 and Bonferroni correction *P* ≤ 0.05), while in hyperglycemia, *MIC26* KO had 3,089 significantly differentially expressed genes (DEGs) as compared to WT-N cells. A clustering analysis of identified transcripts involving all four conditions along with respective replicates is depicted (**Fig S1A**). A Treemap representation shows comparison of significantly upregulated clustered pathways in *MIC26* KOs cultured in normoglycemia compared to WT (**Fig 2A**). Interestingly, the pathways relating to sterol, cholesterol biosynthetic processes and regulation of lipid metabolic processes were significantly upregulated in *MIC26* KO-N compared to WT-N. On the contrary, in *MIC26* KO-H compared to WT-H, pathways involved in sterol, secondary alcohol biosynthetic processes along with cholesterol biosynthesis and cellular amino acid catabolic processes including fatty acid oxidation (FAO) were mainly downregulated (**Fig 2B**). Thus, an antagonistic regulation is observed upon *MIC26* deletion when normoglycemia and hyperglycemia are compared. A detailed pathway enrichment analysis for significantly upregulated genes in *MIC26* KO vs WT cells grown in normoglycemia also revealed genes involved in cholesterol, steroid biosynthetic pathways, fatty acid synthesis and oxidation as well as glycolysis and gluconeogenesis (**Fig 2C**). The genes involved in cholesterol biosynthetic pathways, glycolysis and gluconeogenesis, FAO and fatty acid synthesis were significantly downregulated in *MIC26* KOs grown in hyperglycemia compared to WT cells (**Fig 2D**). The antagonistic behaviour of cholesterol metabolism observed using transcriptomics data (**Fig 2**) was also confirmed in the pathway enrichment analysis for proteomics (**Fig S1B & C**). Detailed analysis of the transcriptomics data in *MIC26* KO-N compared to WT-N showed that ≈80% of the genes regulating cholesterol biosynthesis were significantly upregulated upon normoglycemia (**Fig S2A**) while the opposite was true for hyperglycemia (**Fig S2B**). At the proteome level, the effect of *MIC26* deletion was mainly observed in normoglycemic conditions where 8 out of 12 detected proteins involved in cholesterol biosynthesis showed a significant increase in peptide abundances, while this increase was diminished in *MIC26* KO-H compared to WT-H cells (**Fig S2C-N**). Thus, the loss of MIC26 strongly impacts cholesterol biosynthesis in a nutrient-dependent manner. In conclusion, under normoglycemic conditions, MIC26 acts as a repressor of cholesterol biosynthesis whereas under hyperglycemic conditions MIC26 rather drives this pathway. We further employed targeted metabolomics to decipher any altered cholesterol biosynthesis by quantifying the cholesterol amounts at steady state. In accordance with cholesterol synthesis promoting role of MIC26 in hyperglycemia, cholesterol levels were strongly reduced in *MIC26* KO-H compared to WT-H cells. Moreover, cholesterol levels were significantly increased in WT-H cells compared to WT-N which was not the case and even reversed in *MIC26* KO cells (**Fig S2O**). Thus, MIC26 is required to maintain cholesterol homeostasis and cellular cholesterol demand in a nutrient dependent manner and is of particular importance under hyperglycemia.

**Figure 2.**
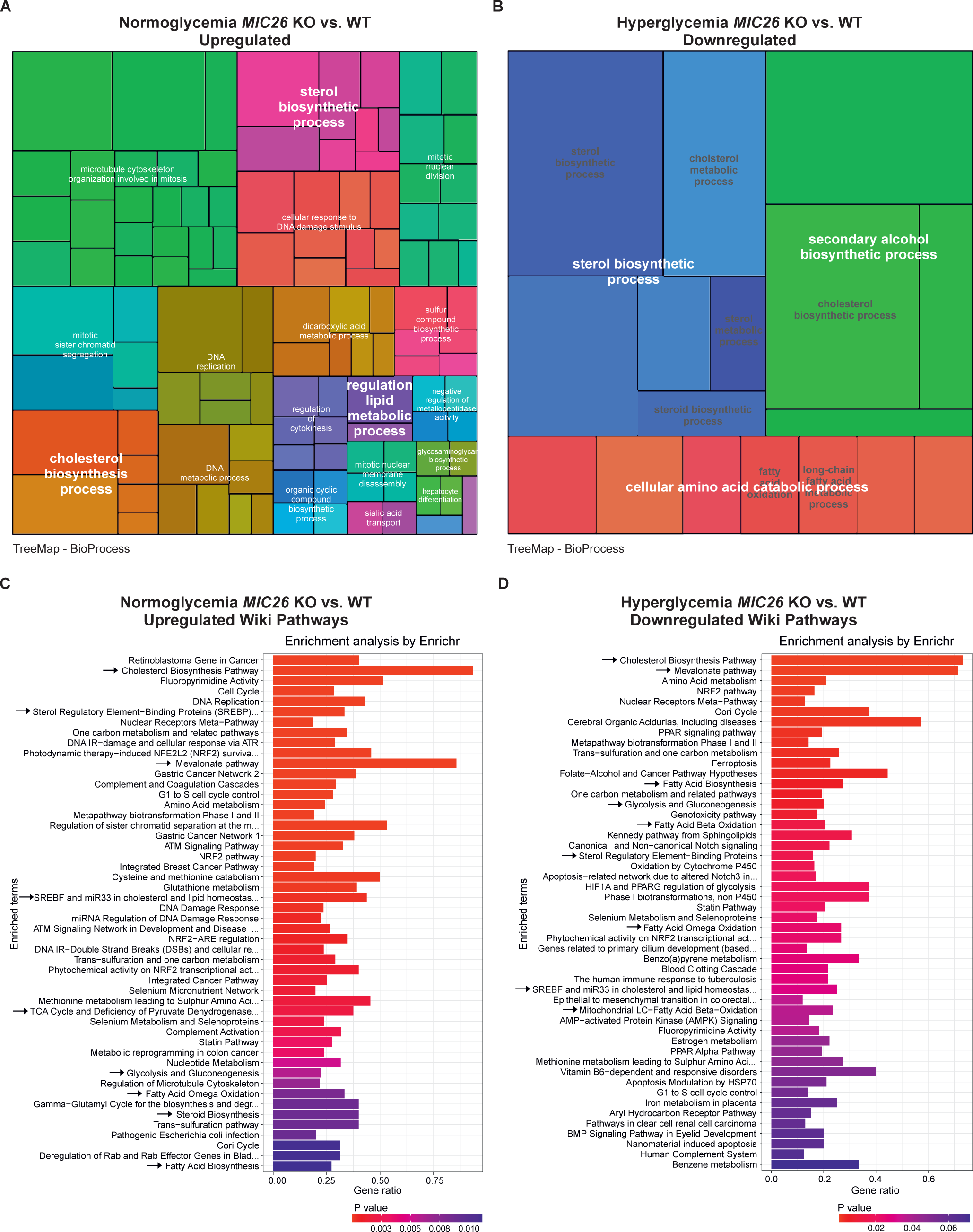
Hyperglycemia confers antagonistic regulation of lipid and cholesterol pathways, in *MIC26* KO vs WT cells, compared to normoglycemia. (A and B) Hierarchical Treemap clustering of significant gene ontology (GO) enriched terms of biological processes upregulated in normoglycemic *MIC26* KO (A) and downregulated in hyperglycemic *MIC26* KO (B) compared to respective WT. Each rectangle represents one BioProcess pathway. Every colour represents clustering of different sub-pathways to pathway families. The rectangle sizes indicate the *P-*value of the respective GO term. (C and D) WikiPathway enrichment using EnrichR analysis of differentially expressed genes (C) upregulated in normoglycemic *MIC26* KO and (D) downregulated in hyperglycemic *MIC26* KO cells compared to respective WT. Arrows indicate antagonistically regulated metabolic pathways including glycolysis, cholesterol biosynthesis, fatty acid synthesis and oxidation. Differentially expressed genes were considered statistically significant with a cut-off fold change of ±1.5 and Bonferroni correction *P* ≤ 0.05. Treemap representation of GO enrichment was plotted with statistically significant pathways with cut-off *P* ≤ 0.05.

### MIC26 maintains the glycolytic function

Besides an antagonistic regulation of the cholesterol biosynthetic pathway, we also observed an opposing trend of genes involved in lipid metabolism as well as glycolysis (**Fig 2C** **& D**). In order to gain further insights about the role of MIC26 regarding the differential regulation of glycolytic pathways in normoglycemia and hyperglycemia, we re-visited our transcriptomics (**Fig S3**) and proteomics (**Fig 3A-C** **and H-J**) datasets and investigated the genes regulating glycolysis upon *MIC26* deletion. On the one hand, in *MIC26* KO-N compared to WT-N, we found that the transcripts encoding hexokinase (*HK*) 1, phosphofructokinase 1 (*PFK1*) (**Fig S3A**) and aldolase (ALDOC) protein levels were significantly upregulated (**Fig 3A**), while glyceraldehyde-3-phosphate dehydrogenase (GAPDH) (**Fig 3B**) and enolase (ENO) were downregulated (**Fig S3A**). On the other hand, in *MIC26* KO-H compared to WT-H, we observed decreased GAPDH and glucose-6-phosphate isomerase (GPI) proteins and transcripts (**Fig 3B** **& C, Fig S3B**). This could indicate that in hyperglycemia, deletion of *MIC26* leads to deregulation of the glycolysis pathway resulting in increased accumulation of glucose (**Fig 3D**) and decreased glycolysis end products. Therefore, to evaluate the metabolic effect of differentially expressed genes (**Fig S3A & B**) and proteins involved in glucose uptake (**Fig 3** **H-J**) and glycolysis (**Fig 3A-C**), we checked whether the glycolytic function is altered in *MIC26* KOs using a Seahorse Flux Analyzer with the glycolysis stress test (**Fig 3E** **& F**). Based on the extracellular acidification rate (ECAR), the ‘glycolytic reserve’ is an index of the ability to undergo a metabolic switch to glycolysis achieved by the cells upon inhibition of mitochondrial ATP generation whereas the ‘glycolytic capacity’ measures the maximum rates of glycolysis which the cell is capable to undergo. Overall ‘glycolytic function’ is measured after cellular glucose deprivation for 1 h and subsequently by quantifying the ECAR primarily arising from cellular lactate formation after providing the cell with saturating glucose amounts. We observed that the glycolytic reserve was significantly increased only in cells cultured in normoglycemia and not in hyperglycemia upon deletion of *MIC26* (**Fig 3F**), while the glycolysis function as well as glycolytic capacity were not significantly increased in *MIC26* KO under both nutrient conditions (**Fig S3C & D**). Therefore, the ability of *MIC26* KO cells (compared to WT) to respond to energetic demand by boosting glycolysis is increased under normoglycemia, while *MIC26* KO cells primed to hyperglycemia were not able to increase glycolytic reserve indicating a clearly different regulation of glycolysis under normoglycemia versus hyperglycemia. In order to understand this better, we quantified the intracellular glucose levels, at steady state in WT and *MIC26* KO, which were significantly increased upon *MIC26* deletion only in hyperglycemia but not normoglycemia compared to the respective WT cells (**Fig 3D**). We further checked whether the increased glucose levels in cells cultured in hyperglycemia is due to increased glucose uptake. In normoglycemia, a glucose uptake assay showed a modest but significant increase of glucose uptake in *MIC26* KOs compared to WT cells (**Fig 3G**) which is consistent with a strong increase in GLUT3 amounts (**Fig 3H**) albeit accompanied by a downregulation of GLUT1 upon *MIC26* depletion (**Fig 3I**). GLUT2 levels remained unchanged in all conditions (**Fig 3J**). However, the observed increased glucose uptake in *MIC26* KO-N (compared to WT-N) was abolished in *MIC26* KO-H (compared to WT-H) and accordingly accompanied by no increase in GLUT3 levels showing that the high amounts of glucose in *MIC26* KO cells grown in hyperglycemia cannot be explained by an increased glucose uptake under these conditions (**Fig 3G**). In *MIC26* KO-N compared to WT-N, even though we observed an increase of glucose uptake, the amount of glycolysis end products, namely pyruvate (**Fig 3K**) and lactate (**Fig 3L**) were unchanged. In hyperglycemia, a significant reduction of pyruvate (**Fig 3K**) and lactate (**Fig 3L**) amounts were observed at steady-state despite increased glucose levels upon *MIC26* deletion. Overall, upon *MIC26* deletion pyruvate and lactate levels were decreased in hyperglycemia while no change was observed in normoglycemia. These results combined with the already discussed differentially regulated transcripts and proteins involved in glycolysis prompted us to check whether there is a difference of shuttling metabolic intermediates from glycolysis towards lipid anabolism. Glycerol-3-phosphate (G3P) is a precursor for lipid biosynthesis synthesized from dihydroxyacetone phosphate which is derived from glycolysis. We observed an increase in G3P levels upon *MIC26* deletion in normoglycemic conditions, while G3P levels were significantly reduced in *MIC26* KO-H cells, compared to the respective WT cells (**Fig 3M**). This opposing trend, together with the previously described antagonistic enrichment in fatty acid biosynthesis (**Fig 2C** **& D**), indicates that *MIC26* deletion rewires glycolytic function to drive lipogenesis in normoglycemia with an antagonistic effect in hyperglycemia. Further, we checked the cellular effect of *MIC26* loss on lipid anabolism in normo-as well as hyperglycemia.

**Figure 3.**
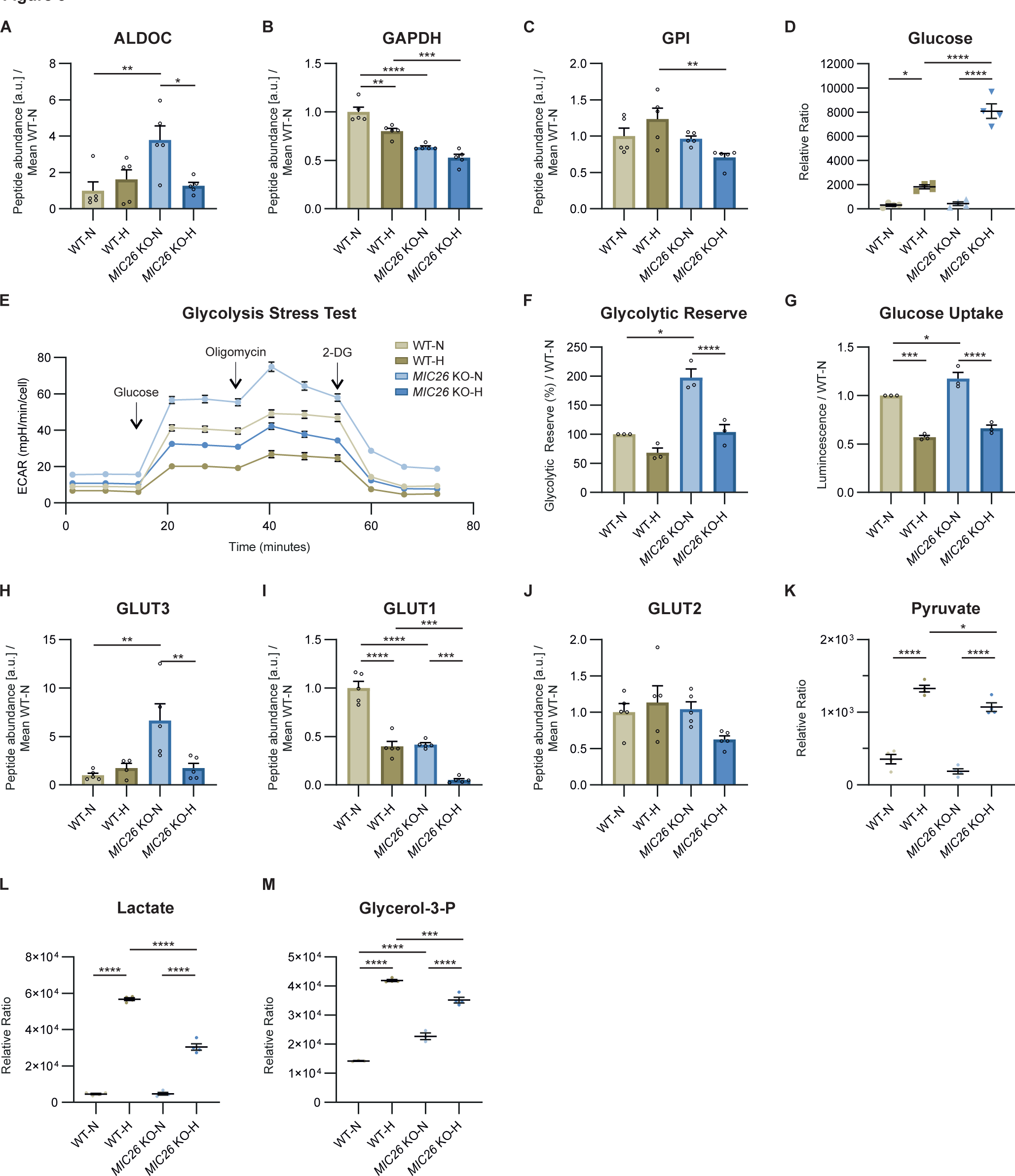
MIC26 maintains the glycolytic function. (A – C) Peptide abundances of enzymes involved in glycolysis pathway curated from proteomics data (N = 5). (D) Steady state metabolomics (GC-MS) data reveals increased cellular glucose accumulation upon *MIC26* deletion in hyperglycemia (N = 3-4). (E and F) Representative glycolysis stress test seahorse assay analysis, with sequential injection of glucose, oligomycin and 2-deoxyglucose, reveals a tendency towards increased glycolysis upon *MIC26* deletion (E) (n = 23). Quantification from various biological replicates shows a significant increase of cellular glycolytic reserve in normoglycemic, but not in hyperglycemic conditions (F) (N = 3). (G) Cellular glucose uptake was measured using Glucose uptake Glo assay normalized to WT-N. *MIC26* deletion leads to an increased glucose uptake upon normoglycemia (N = 3). (H – J) Peptide abundances of transporters involved in glucose uptake namely GLUT3 (H), GLUT1 (I) and GLUT2 (J) curated from proteomics data (N = 5). (K and L) Steady state metabolomics (GC-MS) shows unaltered cellular pyruvate (K) and lactate (L) levels in *MIC26* KO cell lines in normoglycemia but decreased levels upon *MIC26* deletion in hyperglycemia (N = 3-4). (M) *MIC26* deletion increases glycerol-3-phosphate amount in normoglycemia with an antagonistic effect in hyperglycemia compared to the respective WT (N = 3-4). Data are represented as mean ± SEM (A-M). Statistical analysis was performed using one-way ANOVA with \**P* < 0.05, \*\**P* < 0.01, \*\*\**P* < 0.001, \*\*\*\**P* < 0.0001. N represents the number of biological replicates and n the number of technical replicates.

### The loss of MIC26 leads to metabolic rewiring of cellular lipid metabolism via CPT1 and dysregulation of fatty acid synthesis

The respective increase and decrease of G3P (**Fig 3M**) in normoglycemia and hyperglycemia upon *MIC26* deletion when compared to WT as well as an opposing trend in fatty acid biosynthesis reflected in our transcriptomics data (**Fig 2C** **& D**) prompted us to explore the regulation of cellular lipid metabolism. Lipid droplets (LDs) play a key role in energy metabolism and membrane biology by acting as reservoirs to store TAG and sterol esters which are released to the relevant pathways according to cellular demand (Thiam *et al*, 2013). Using BODIPY staining, we checked the cellular LD content in unstimulated and palmitate-stimulated WT and *MIC26* KO cells grown in normoglycemia and hyperglycemia, respectively. The number of LDs and the respective fluorescence intensity of BODIPY are indicative of cellular lipid content (Chen *et al*, 2022). We observed a general increase of LD number in *MIC26* KOs irrespective of treatment conditions (**Fig 4A** **& B**). However, the increased intensity of BODIPY staining observed in normoglycemia was not evident in *MIC26* KO-H compared to respective WT cells (**Fig 4C** **& D**). Further, when we fed free fatty acids (FFAs) in the form of palmitate, there was again increased BODIPY intensity in *MIC26* Kos in normoglycemia even at a higher level. In contrast, under hyperglycemia *MIC26* KO cells showed lower LD intensity when compared to WT cells again demonstrating an antagonistic role of MIC26 when normoglycemia was compared to hyperglycemia (**Fig 4C** **& D**). These experiments allow us to conclude that the effect of *MIC26* deletion on LD accumulation depends on the nutrient condition which is enhanced under nutrient-rich (high-glucose/high-fat-like) conditions. Overall, MIC26 is essential to regulate the amount of cellular LD content in a nutrient-dependent manner (**Fig 4D**).

**Figure 4.**
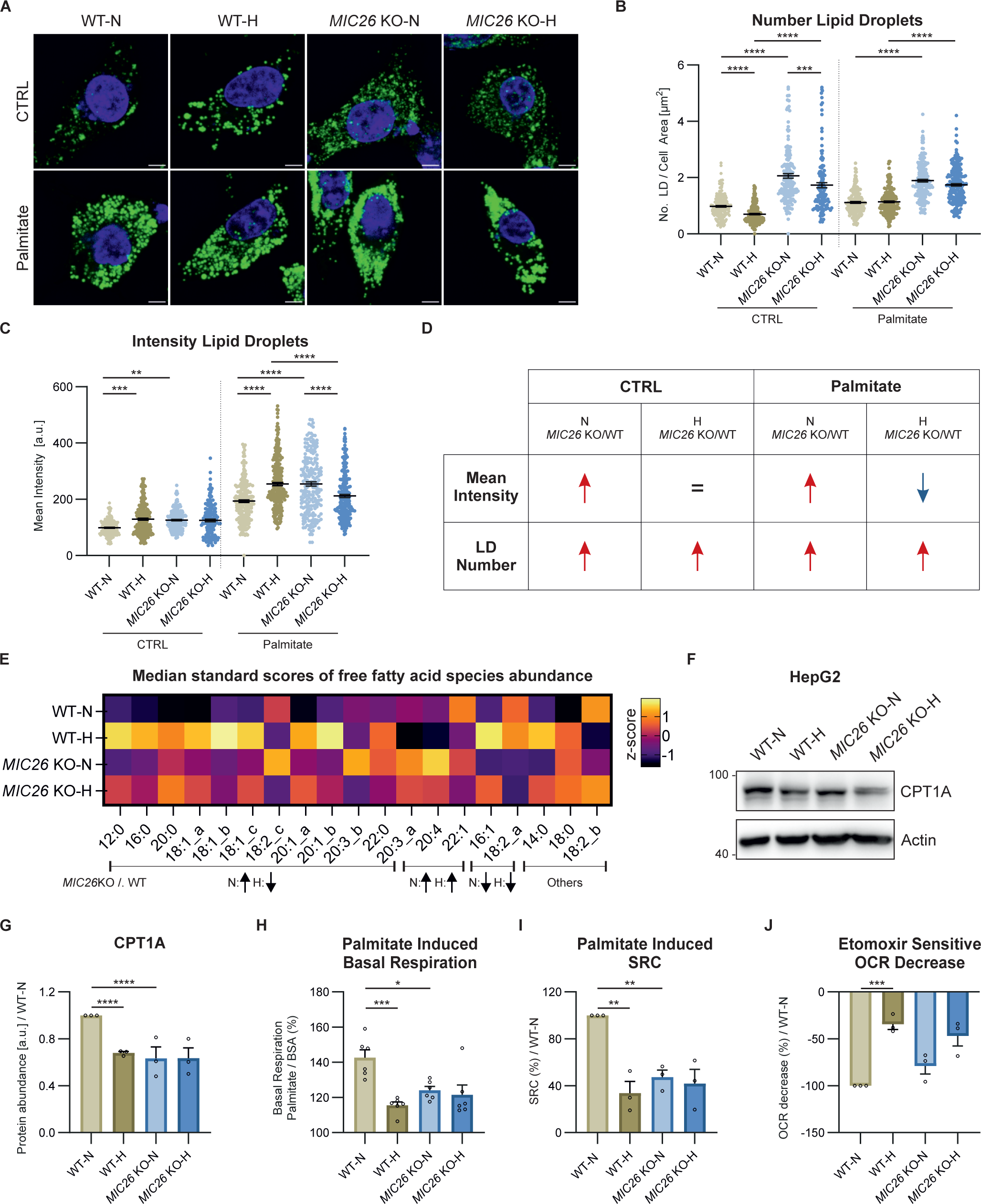
The loss of MIC26 leads to metabolic rewiring of cellular lipid metabolism via CPT1A and dysregulation of fatty acid synthesis. (A – D) Analysis of lipid droplet formation in WT and *MIC26* KO cells cultured in normo- and hyperglycemia either in standard growth condition (CTRL) or upon palmitate stimulation (100 µM, 24 h). Representative confocal images of lipid droplets stained using BODIPY 493/503 are shown (A). Quantification shows number of lipid droplets normalized to the total cell area [µm^2^] (B) and mean fluorescence intensity per cell normalized to mean intensity of WT-N in all biological replicates (C). *MIC26* deletion leads to a nutritional-independent increase in lipid droplet number. However, an opposing effect, leading to increase or decrease of mean fluorescence intensity of lipid droplets, upon comparison of *MIC26* KO to WT was observed in normo- and hyperglycemia respectively, with a pronounced effect upon feeding palmitate (N = 3). Scale bar represents 5 µm. (E) Heat map representing the abundance of steady state FFA species in WT and *MIC26* KO cells cultured in normo- and hyperglycemia. 11 out of 19 of the FFA species represent an antagonistic behavior upon comparing *MIC26* KO to WT in normo- (increase) and hyperglycemia (decrease) (N = 3-4). (F and G) Western blot analysis (F), along with respective quantification (G) of WT and *MIC26* KO cells cultured in normo- and hyperglycemia, show a reduction of CPT1A in WT-H, *MIC26* KO-N and *MIC26* KO-H compared to WT-N (N = 3). (H – J) Mitochondrial fatty acid oxidation analyzed using Seahorse XF analyzer shows a decreased palmitate-induced basal respiration (H) and spare respiratory capacity (I) and a nonsignificant reduction of etomoxir-sensitive OCR decrease upon comparing *MIC26* KO to WT in normoglycemia (N = 3). Data are represented as mean ± SEM (B-C and G-J). Statistical analysis was performed using one-way ANOVA with \**P* < 0.05, \*\**P* < 0.01, \*\*\**P* < 0.001, \*\*\*\**P* < 0.0001. N represents the number of biological replicates.

LD biogenesis is closely linked to increased cellular FFA levels (Zadoorian *et al*, 2023). Using targeted metabolomics, we investigated the steady state levels of long chain FFAs in WT and *MIC26* KO cell lines cultured in normoglycemia and hyperglycemia (**Fig 4E**). We identified that there was either no change or an increase of saturated FFAs including lauric (12:0), myristic (14:0), palmitic (16:0), stearic (18:0), arachidic (20:0) as well as behenic (22:0) acid in normoglycemia in *MIC26* KOs compared to WT cells (**Fig 4E, Fig S4**). In contrast, we consistently found a decrease in most of the above-mentioned saturated FFAs in *MIC26* KO-H compared to WT-H. This trend was also observed in unsaturated FFAs like oleic acid (18:1). Overall, we conclude that there is a consistent decrease of saturated FFAs in *MIC26* KO-H as opposed to *MIC26* KOs grown in normoglycemic conditions consistent to the observed trend in LD formation. Increased level of FFAs and LDs can arise from increased FFAs biosynthesis as well as reduced FFA catabolism via mitochondrial β-oxidation (Afshinnia *et al*, 2018). Mitochondrial β-oxidation requires import of long chain FFAs using the carnitine shuttle comprised of carnitine palmitoyl transferase 1 (CPT1) and 2 (CPT2) and carnitine-acylcarnitine translocase (CACT), into the mitochondrial matrix. Depletion of CPT1A, which is the rate limiting step of FAO, coincides with lipid accumulation in the liver (Sun *et al*, 2021). Therefore, we determined the CPT1A amounts using WB analysis (**Fig 4F** **& G**) which were in line with transcriptomics and quantitative PCR data (**Fig S5A & B**). *MIC26* deletion revealed a reduction of CPT1A in normoglycemia compared to WT cells. In WT cells, hyperglycemia already triggered a reduction in CPT1A level and there was no further decrease of CPT1A in *MIC26* KOs grown in hyperglycemia (**Fig 4F** **& G**). In order to understand the functional significance of CPT1A reduction on mitochondrial function, we checked the FAO capacity of respective cell lines by feeding them with palmitate and analysing the induced basal respiration and spare respiratory capacity (SRC) of mitochondria compared to BSA control group (**Fig 4H and I, Fig S5C & D**). SRC is the difference between FCCP stimulated maximal respiration and basal oxygen consumption and therefore is the ability of the cell to respond to an increase in energy demand. We observed a significant reduction in palmitate-induced basal respiration as well as SRC in *MIC26* KO-N compared to WT-N determining decreased mitochondrial long chain fatty acid β-oxidation. It is important to note that we already observed a significant decrease in mitochondrial β-oxidation in WT-H condition which was not further affected in *MIC26* KO-H in agreement with the reduced CPT1A levels (**Fig 4F** **& G**). We further analysed the reduction of oxygen consumption rate (OCR) induced by etomoxir inhibition of CPT1A (**Fig 4J**). In *MIC26* KO-N compared to WT-N, palmitate-induced OCR was reduced moderately, yet this was not significant. For the respective hyperglycemic conditions, we did not observe a change which was again in line with the observed CPT1A levels. Thus, reduced β-oxidation in *MIC26* KO-N compared to WT-N is apparently contributing to increased FFA levels and LD content and could be mediated, at least in part, via the reduced levels of CPT1A resulting in reduced transport of FFAs into mitochondria.

We further checked whether FFA biosynthesis plays a role in the nutrition-dependent antagonistic regulation of lipid anabolism in *MIC26* KO cell line. FFA biosynthesis is initiated with the export of citrate generated in TCA cycle from mitochondria to the cytosol. The export is mediated by the citrate/malate exchanger SLC25A1 which is present in the mitochondrial IM. Proteomics and transcriptomics data showed that SLC25A1 was increased in normoglycemia in *MIC26-*KOs (compared to respective WT), but not in hyperglycemia (**Fig S5E & F**). We then checked for further changes in the transcriptome and proteome levels of key enzymes playing a role in FFA synthesis. We found that ATP citrate lyase (ACLY, **Fig S5G**), acetyl-Co-A carboxylase (ACACA, **Fig S5H &I**) which converts acetyl-CoA into malonyl-CoA, fatty acid synthase (FASN, **Fig S5J & K**) and acetyl-CoA desaturase (SCD, **Fig S5L & M**) were increased in normoglycemia in *MIC26* KOs but mostly unchanged in hyperglycemia. In addition, hyperglycemia resulted in an increase of glycerol kinase (GK) in WT cells which was absent in *MIC26* KO cells (**Fig S5N**). Therefore, our data indicate that the FFA biosynthesis pathway is upregulated upon loss of *MIC26* KO in normoglycemia but not in hyperglycemia compared to respective WT conditions. An upregulation of FFA biosynthesis along with reduced mitochondrial β-oxidation partially mediated by reduced CPT1A amount in *MIC26* KO-N and a shift of glycolytic intermediates resulting in G3P accumulation show that loss of MIC26 leads to a cumulative metabolic rewiring towards increased cellular lipogenesis.

### *MIC26* deletion leads to hyperglycemia-induced decrease in TCA cycle intermediates

To synthesize FFA, citrate first needs to be generated by the TCA cycle in the mitochondrial matrix before it is exported to the cytosol. Using targeted metabolomics, we checked whether the TCA cycle metabolism is altered upon *MIC26* deletion at steady state in both nutrient conditions (**Fig 5A**). As previously described, glycolysis resulted in decreased pyruvate levels upon *MIC26* deletion in hyperglycemia, while no change was observed in normoglycemia (**Fig 3N**). Furthermore, most of the downstream metabolites including (iso-)citrate, succinate, fumarate and malate consistently showed a significant decrease in *MIC26* KO cells cultured in hyperglycemia, compared to WT condition, but not in normoglycemia following the previously observed trend in pyruvate levels. To elucidate a possible defect of mitochondrial pyruvate import, we checked mitochondrial pyruvate carrier 1 (MPC1) and MPC2 abundances (**Fig 5B** **& C**) as well as mitochondrial respiration after blocking mitochondrial pyruvate carrier (glucose / pyruvate dependency) using UK5099 inhibitor (**Fig 5D**). While we observed a downregulation of MPC1 in *MIC26* KO-N compared to WT-N, MPC1 abundances in *MIC26* KO-H compared to WT-H remained unchanged. Further we did not observe any changes in MPC2 level. Also, mitochondrial glucose/pyruvate dependency remained unchanged in the respective hyperglycemia combination while we observed a minor but significant decrease in *MIC26* KO-N compared to WT-N. In addition, elucidation of abundances of mitochondrial enzymes catalyzing TCA cycle metabolites (**Fig S6A-K**) as well as the respective cytosolic enzymes (**Fig S6L-N**) interestingly revealed an upregulation of citrate synthase (**Fig S6A**) and mitochondrial aconitase 2 (**Fig S6B**) in *MIC26* KO cells independent of nutrient conditions. Furthermore, the immediate downstream enzyme isocitrate dehydrogenase 2 which generates α-ketoglutarate (α-KG) was upregulated in *MIC26* KO condition (**Fig S6C**). In contrast to all previously described metabolites, the α-KG levels were increased in hyperglycemia in *MIC26* KO compared to WT. The accumulation of α-KG possibly arises from a significant downregulation of α-KG dehydrogenase in *MIC26* KO independent of the nutrient condition (**Fig S6E**). Following, an accumulation of α-KG by downregulation of α-KG dehydrogenase would further explain the decreased formation of succinate in *MIC26* KO-H compared to WT-H. Succinate dehydrogenases (**Fig S6G & H**) as well as fumarase (**Fig S6I**) did not show any changes in abundances upon the respective *MIC26* KO to WT comparison reflecting the uniform metabolite trend in succinate, fumarate and malate. Overall, we observed a general decrease in several TCA cycle metabolites in *MIC26* KO-H compared to WT-H. Therefore, we propose that a downregulation of FFA biosynthesis in *MIC26* KO-H compared to WT-H results from a limited formation of citrate via the mitochondrial TCA cycle presumably arising from reduced utilization of glucose.

**Figure 5.**
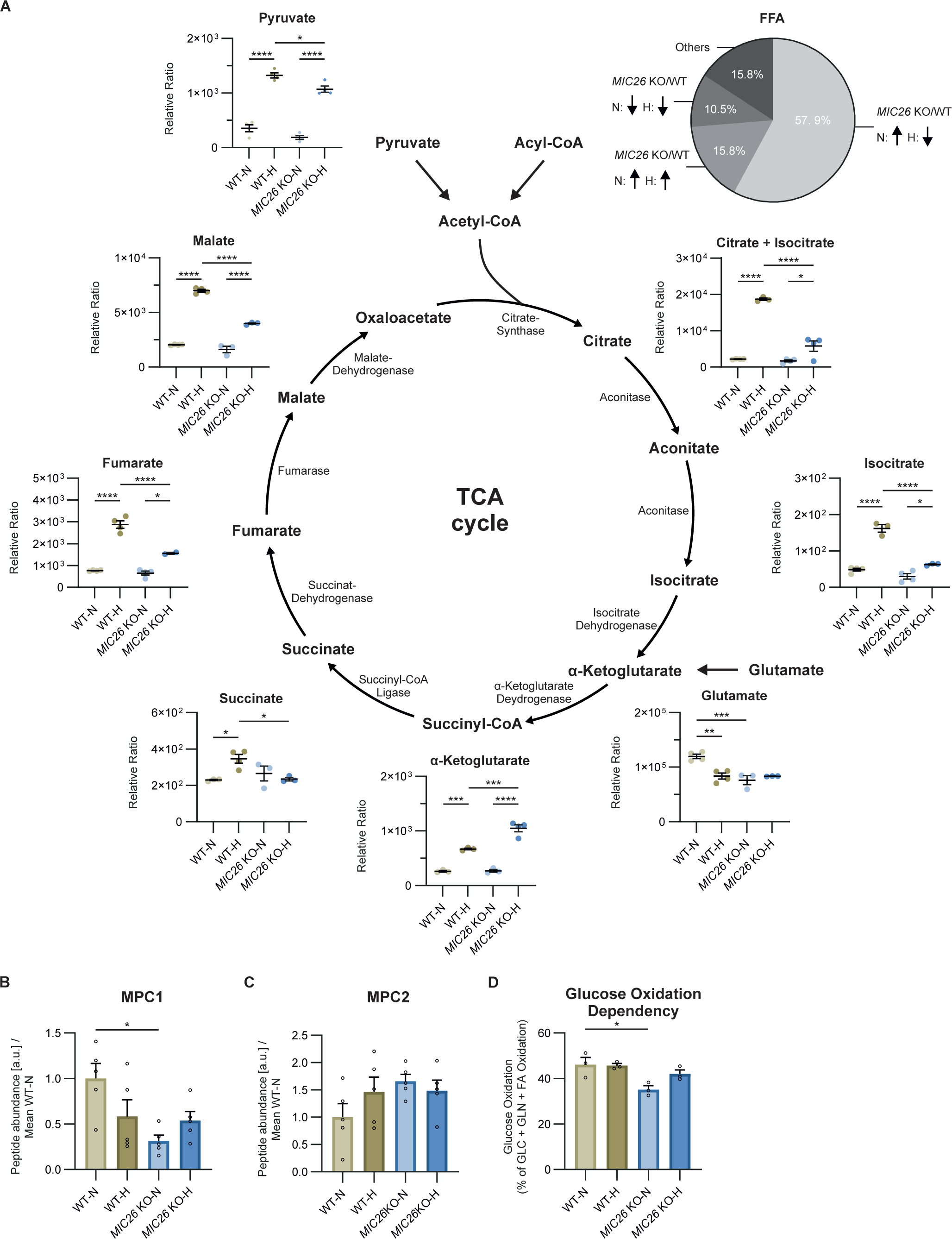
*MIC26* deletion leads to hyperglycemia-induced decrease in TCA cycle intermediates. (A) Representation of the relative amounts (GC-MS) of TCA cycle metabolites and associated precursors at steady state in WT and *MIC26* KO cells cultured in normo- and hyperglycemia. All the TCA cycle metabolites with the exception of α-ketoglutarate showed a decreasing trend upon *MIC26* KO when compared to WT in hyperglycemia (N = 3-4). (B and C) Mitochondrial pyruvate carrier 1 (MPC1) (B), but not MPC2 (C), is significantly decreased in *MIC26* KO-N compared to WT-N, as revealed by peptide abundances from proteomics data (N = 5). (D) Mitochondrial glucose / pyruvate dependency analysis, using Seahorse XF analyzer mito fuel flex test assay, reveals a decreased mitochondrial respiratory dependency of *MIC26* KO on glucose / pyruvate in normoglycemia (N = 3). Data are represented as mean ± SEM (A-C). Statistical analysis was performed using one-way ANOVA with \**P* < 0.05, \*\**P* < 0.01, \*\*\**P* < 0.001, \*\*\*\**P* < 0.0001. N represents the number of biological replicates.

### Aberrant glutamine metabolism is observed in *MIC26* KOs independent of nutritional status

Glutaminolysis feeds α-KG in the TCA cycle. To check whether the increase in α-KG could be (apart from downregulation of α-KG dehydrogenase amounts) derived from glutaminolysis, we also checked glutamine (**Fig 6A**) and glutamate levels (**Fig 5A**). The amounts of glutamine at steady-state were uniformly increased in *MIC26* KOs irrespective of nutrient conditions (**Fig 6A**). Glutamate was decreased in *MIC26* KO-N compared to WT-N (**Fig 5A**). Mitochondria mainly oxidise three types of cellular fuels namely pyruvate (from glycolysis), glutamate (from glutaminolysis) and FFAs. We used a ‘mito-fuel-flex-test’ for determining the contribution of glutamine as a cellular fuel. The contribution of glutamine as cellular fuel could be determined using BPTES, an allosteric inhibitor of glutaminase (GLS1), which converts glutamine to glutamate. The extent of reduction of mitochondrial oxygen consumption upon BPTES inhibition is used as a measure for determining the glutamine dependency while the capacity is the ability of mitochondria to oxidise glutamine when glycolysis and FFA oxidation are inhibited. Intriguingly, we observed that the *MIC26* KOs do not depend on glutamine as a fuel (**Fig 6B**, left histogram). However, they still can use glutamine when the other two pathways were inhibited (**Fig 6B**, right histogram). The glutamine oxidation capacity of *MIC26* KO cells cultured in normoglycemia as well as hyperglycemia appears slightly decreased compared to WT but this decrease is not statistically significant. Overall, we observe a remarkable metabolic rewiring of *MIC26* KOs to bypass glutaminolysis. In order to understand whether the independency on glutamine as fuel arises due to the possibility of aberrant transport of glutamine into the mitochondria, we analysed transcripts and proteins that were not only significantly downregulated but also present in the mitochondria IM and interacted with MIC26. For this, we investigated putative MIC26 interactors by compiling a list using BioGRID, NeXtProt and IntAct databases. SLC25A12, an antiporter of cytoplasmic glutamate and mitochondrial aspartate, was significantly downregulated (**Fig S7A**) while showing up in the interactome of MIC26 (**Fig S7E**). Accordingly, WB analysis reveal a reduction of SLC25A12 in *MIC26* KOs compared to WT HepG2 cells in both normoglycemia and hyperglycemia (**Fig 6C** **& D**). Further, it is known that a variant of SLC1A5 transcribed from an alternative transcription start site and present in the mitochondrial IM is responsible for transporting glutamine into mitochondria (Yoo *et al*, 2020a). Transcriptomics data revealed a reduction of SLC1A5 in *MIC26* KOs while proteomics revealed a significant reduction in normoglycemia and non-significant reduction in hyperglycemia in *MIC26* KOs compared to WT (**Fig S7B & C**). An increase of cellular glutamine levels in *MIC26* KOs (**Fig 6A**) along with reduced levels of SLC1A5 and reduced mitochondrial glutamine dependency (**Fig 6B**) indicate a reduced transport of glutamine destined for glutaminolysis into mitochondria.

**Figure 6.**
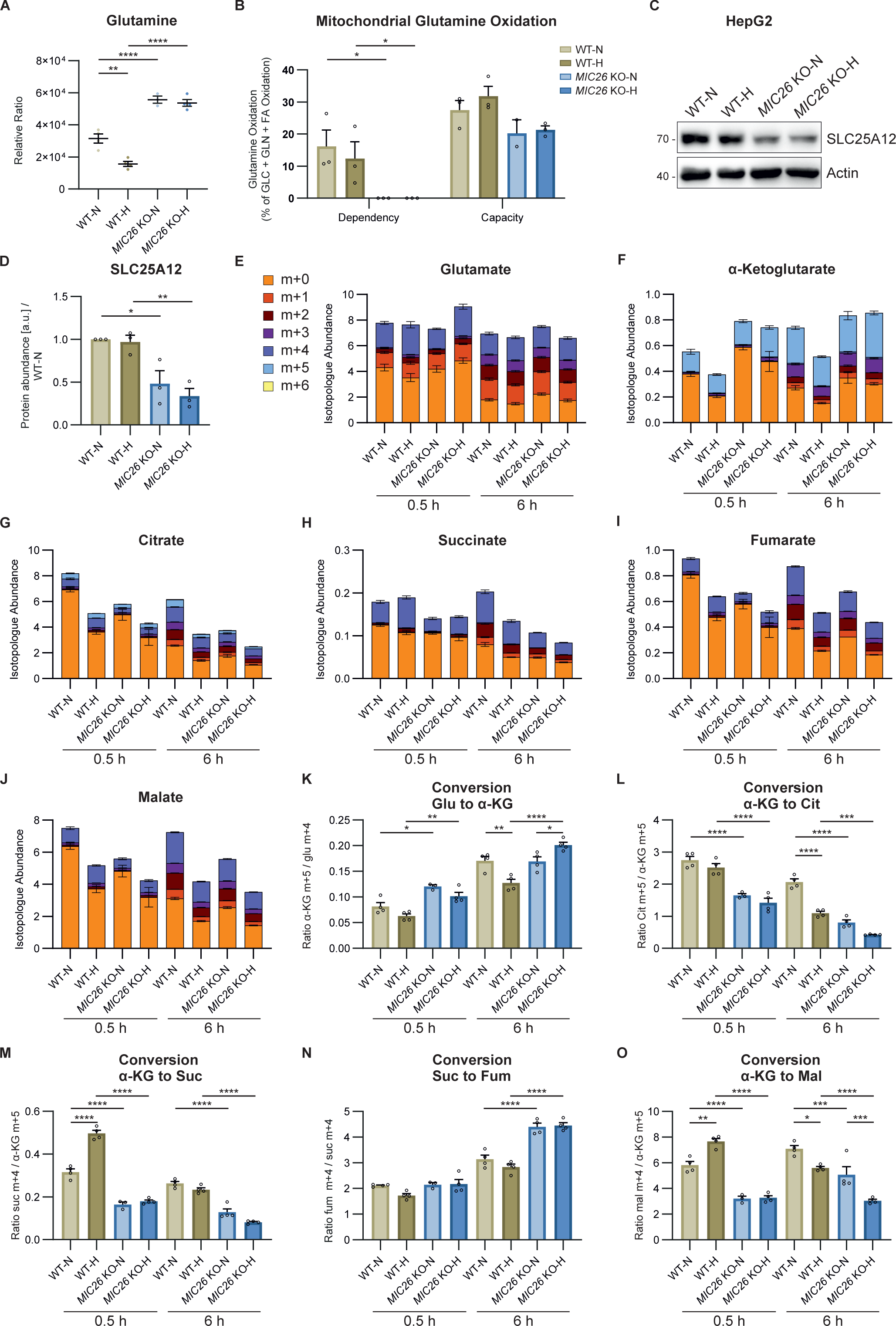
Aberrant glutamine metabolism is observed in *MIC26* KOs independent of nutritional status. (A) Metabolomics analysis (GC-MS) shows that glutamine levels were strongly increased in *MIC26* KO cells cultured in both normo- and hyperglycemia at steady state compared to respective WT (N = 3-4). (B) Quantification of mitochondrial glutamine dependency and capacity analysis, using Seahorse XF analyzer mito fuel flex test assay, shows a diminished mitochondrial respiratory dependency on glutamine. A nonsignificant mitochondrial respiratory decreased capacity of *MIC26* KO cells was observed compared to respective WT conditions (N = 3). (C and D) Western Blot analysis (C) along with respective quantification (D) show reduced amounts of the glutamate aspartate antiporter SLC25A12 (ARALAR / AGC1), present in mitochondria, in *MIC26* KO cell lines compared to respective WT cells (N = 3). (E – J) Representation of labeled (m+1 - m+6) and unlabeled (m+0) species of glutamate (GC-MS) (E), and TCA cycle metabolites (AEC-MS) α-KG (F), citrate (G), succinate (H), fumarate (I) and malate (J), from glutamine tracing experiments after labelling for 0.5 h and 6 h (N = 4). (K – O) Conversion rates from different TCA cycle reactions calculated using the ratio of highest labeled species abundances for the conversions of glutamate to α-KG (K), α-KG to citrate (L), α-KG to succinate (M), succinate to fumarate (N) and α-KG to malate (N = 4). Data are represented as mean ± SEM (A-B and D-O). Statistical analysis was performed using one-way ANOVA with \**P* < 0.05, \*\**P* < 0.01, \*\*\**P* < 0.001, \*\*\*\**P* < 0.0001. N represents the number of biological replicates.

In order to delineate whether the increased glutamine levels at steady-state are due to decreased glutamine utilisation or increased flux, we performed a metabolic tracing experiment where WT and *MIC26* KO cells, cultured in normoglycemia and hyperglycemia, were fed with labelled glutamine [U-^13^C_5_, ^15^N_2_] for 0.5 h and 6 h (**Fig 6E-O**). Glutamine is converted to glutamate by glutaminase (GLS) in the mitochondria. The GLS amounts were not altered in *MIC26* KO cells when compared to respective WT cells grown in normoglycemia and hyperglycemia (**Fig S7D**). In line, label enriched glutamate species (m+1 - m+4) did not show major differences in all four conditions at both timepoints (**Fig 6E**). Following this, we hypothesize that accumulation of glutamine in *MIC26* KO cells arises from other cellular pathways utilising glutamine being impaired, for example synthesis of purine, pyrimidine or amino acids. However, labelled α-KG (m+1 - m+5) was increased upon *MIC26* deletion with a pronounced effect in cells cultured in hyperglycemia similar to the detected steady-state amounts of α-KG (**Fig 6F**). To check the conversion rates of different metabolite reactions, we determined the enzyme conversion rates by calculating the ratio of the highest labelled species from the end-metabolite compared to the starting-metabolite. In accordance to the observed level of α-KG, the conversion ratio from glutamate to α-KG was significantly increased in *MIC26* KO cells (**Fig 6K**). We further checked the flux of TCA metabolites downstream to α- KG namely succinate, fumarate and malate. Despite the increased α-KG levels, the labelled succinate species (m+1 - m+4) was decreased in *MIC26* KO cells (**Fig 6H**). In line, the conversion rate from α-KG to succinate was significantly downregulated in *MIC26* KO cells independent of glucose concentrations and timepoints (**Fig 6M**). However, the conversion ratio from succinate to fumarate catalysed by mitochondrial complex II subunits, succinate dehydrogenases A-D, was increased in *MIC26* KO cell lines at the 6 h timepoint compared to WT in both normoglycemia and hyperglycemia (**Fig 6N**). Despite the increase in fumarate conversion, the labelled fumarate and malate were decreased in *MIC26* KO compared to WT in normoglycemia but not in hyperglycemia (**Fig 6I** **& J**) while there were minor differences at 0.5 h. The conversion ratio from α-KG to malate was decreased upon *MIC26* deletion in both nutrient conditions at 0.5 h and 6 h of glutamine labelling. Thus, despite increased conversion of succinate to fumarate as well as increased flux from glutamate to α-KG (in hyperglycemia) upon loss of MIC26, cellular glutaminolysis does not function optimally. We also checked the labelled citrate levels which showed minor changes after 0.5 h treatment but a major change in all labelled species (m+1 - m+5) after 6 h (**Fig 6G**). Correspondingly, the levels of citrate in WT-N cells were highly increased compared to all three other conditions. Conversion rates from α-KG (m+5) to citrate (m+5) were significantly reduced in *MIC26* KO cell lines compared to the respective WT cells (**Fig 6L**). Overall, the flux of glutamine through the TCA cycle is accompanied by decreased conversion of TCA cycle intermediates. Therefore, we conclude that aberrant glutaminolysis is observed upon loss of MIC26.

### MIC26 regulates mitochondrial bioenergetics by restricting the ETC activity and OXPHOS (super-)complex formation

We have shown that the loss of *MIC26* leads to dysregulation of various central fuel pathways. In order to understand the effect of *MIC26* deletion on cellular bioenergetics, we checked the mitochondrial membrane potential (ΔΨ_m_) of WT and *MIC26* KO cells in both nutrient conditions by employing TMRM dye (**Fig 7A** **& B**). Loss of MIC26 leads to decreased ΔΨ_m_ compared to control cells in both normoglycemia and hyperglycemia. It is well known that mitochondrial loss of membrane potential is connected to mitochondrial dynamics (Giacomello *et al*, 2020). Thus, we checked the mitochondrial morphology and observed that loss of MIC26 consistently leads to a significant increase of mitochondrial fragmentation compared to WT-N (**Fig 7C** **& D**). In addition, WT cells grown in hyperglycemia despite maintaining the ΔΨ_m_ exhibited fragmented mitochondria. We also checked the levels of major mitochondrial dynamic regulators: MFN1, MFN2, DRP1 as well as OPA1 processing into short forms. WB analysis showed that MFN1 levels were significantly decreased upon *MIC26* deletion in both normoglycemia and hyperglycemia compared to respective WT cells (**Fig S8A & B**) which could account for increased fragmentation. There was no major effect on the amounts of other factors which could account for mitochondrial fragmentation. Thus, *MIC26* deletion is characterized by reduced ΔΨ_m_ and fragmentation of mitochondria which indicate altered mitochondrial bioenergetics. To determine this, we checked the mitochondrial function in WT and *MIC26* KO cells by using a mitochondrial oxygen consumption assay (**Fig 7E**). We observed an increased basal respiration in *MIC26* KOs in both normoglycemia and hyperglycemia compared to the respective WT (**Fig 7F**). The ATP production was increased in *MIC26* KO cells in hyperglycemia compared to WT-H (**Fig S8C**). In addition, decreased SRC was observed in *MIC26* KO-N when compared to WT-N condition (**Fig S8D**). Overall, *MIC26* KOs demonstrate higher basal respiration in both nutrient conditions. In order to elucidate the increased basal respiration, we performed blue native PAGE to understand the assembly of OXPHOS complexes along with in-gel activity assays (**Fig 7G**). *MIC26* deletion consistently led to an increase in the levels of OXPHOS complexes I, III, IV and dimeric and oligomeric complex V (shown in green arrows) (**Fig 7G**, left blots respectively for each complex). The increased assembly of OXPHOS complexes was also accompanied by respective increase of in-gel activity (shown in blue arrows) (**Fig 7G**, right blots respectively for each complex). This is consistent with the previously observed increased basal respiration (**Fig 7F**) and the succinate to fumarate conversion representing an increased complex II activity (**Fig 6N**). Altogether, we conclude that formation and stability of OXPHOS (super-) complexes as well as their activity is dependent on MIC26.

**Figure 7.**
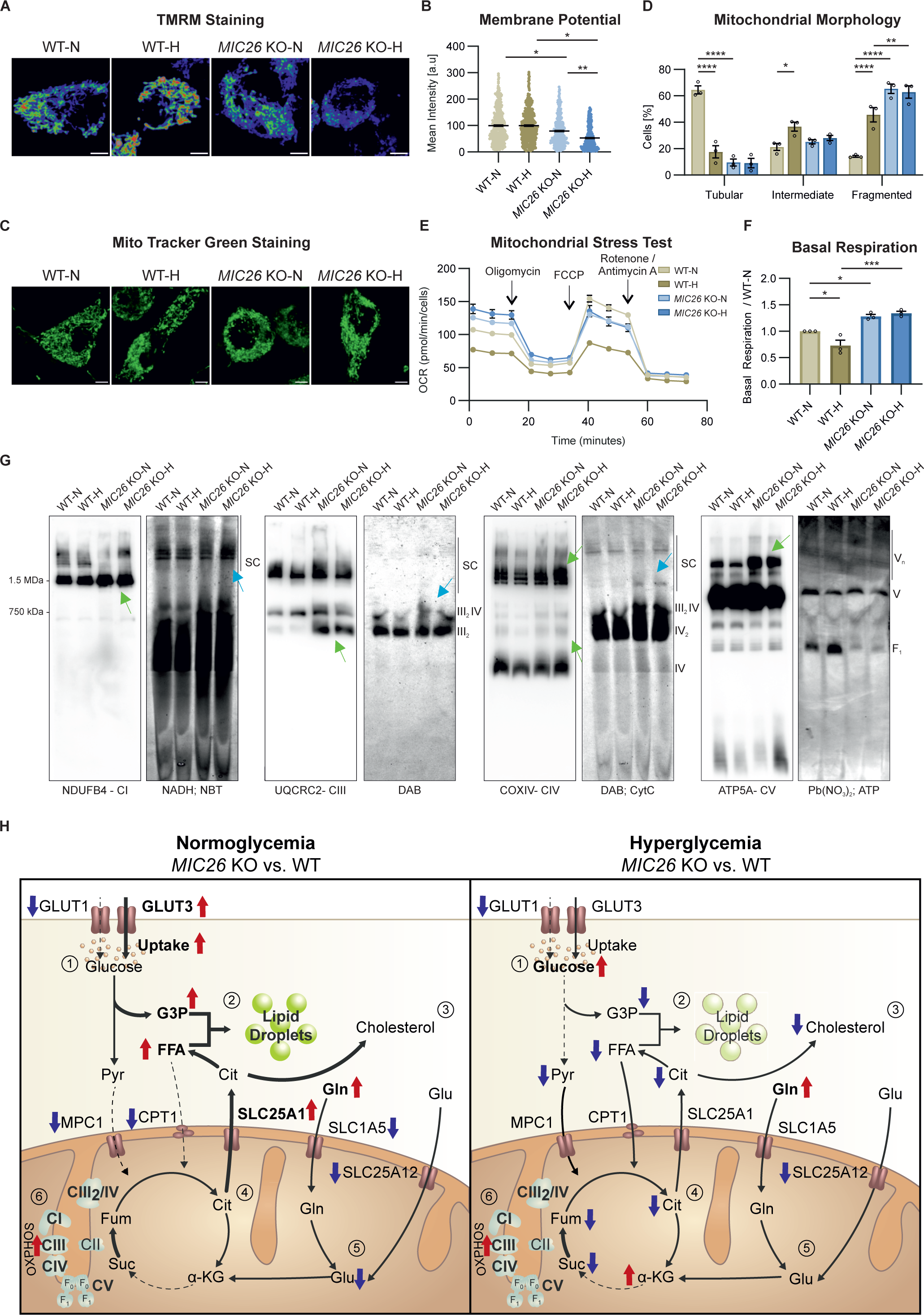
MIC26 regulates mitochondrial bioenergetics by restricting the ETC activity and OXPHOS (super-)complex formation. (A and B) Representative pseudocolour rainbow LUT intensities from confocal images of WT and *MIC26* KO HepG2 cells stained with TMRM show a reduction in ΔΨ_m_ upon *MIC26* deletion in both normoglycemia and hyperglycemia when compared to respective WT cells (A). Quantification represents mean TMRM fluorescence intensity per cell normalized to mean intensity of WT-N in all biological replicates (B) (N = 3). Scale bar represents 5 µm. (C and D) Representative confocal images of mitochondrial morphology, visualized by MitoTracker green staining (C), show that loss of MIC26 shifts mitochondrial morphology from tubular mitochondrial network in WT normoglycemic conditions to fragmented phenotype irrespective of supplemented glucose amount (D) (N = 3). Scale bar represents 5 µm. (E and F) Representative mitochondrial stress test with Seahorse XF analyzer, with sequential injection of oligomycin, FCCP and rotenone/antimycin (E) (n = 19-23). Quantification from various biological replicates shows a significant increase of basal respiration in *MIC26* KOs cultured in both normo- and hyperglycemia (F) (N = 3). (G) Blue native (respective left panel) and clear native (respective right panel) PAGE analysis reveals an overall increase of OXPHOS complex formation (for CI, CIII, CIV and CV, green arrows) as well as corresponding increased in-gel activity of supercomplexes, and complex III_2_IV (blue arrows) upon *MIC26* deletion. CV shows no in-gel activity alterations while a decreased in-gel activity of F_1_ occurs upon loss of MIC26. Native PAGEs were performed in three biological replicates and representative gels are shown. (H) Model representing the antagonistic regulation of metabolic pathways encompassing glucose usage, lipid droplet formation, cholesterol synthesis, as well as decrease in TCA cycle metabolites in MIC26 deficient HepG2 cells dependent on nutritional conditions compared to respective WT cells. An increase of glutamine levels as well as assembly of various OXPHOS complexes is observed in *MIC26* KOs independent of the nutritional status. Arrows indicate respective up (red) or downregulated (blue) protein/metabolite or activity levels, respectively. In the model, left panel indicates normoglycemic while the right panel represents the hyperglycemic conditions. Data are represented as mean ± SEM (B and D-F). Statistical analysis was performed using one-way ANOVA with \**P* < 0.05, \*\**P* < 0.01, \*\*\**P* < 0.001, \*\*\*\**P* < 0.0001. N represents the number of biological replicates and n the number of technical replicates.

## Discussion

Our study identifies MIC26 as a critical regulator at the crossroads of several major metabolic pathways. Based on detailed multi-omics analyses, we deciphered an intricate interplay between MIC26, a mitochondrial IM protein, and global cellular metabolic adaptations. To understand these metabolic changes and their dependency on mitochondrial ultrastructure and function is of high medical relevance as nutrient-overload is known to cause obesity and T2DM in humans. In fact, *MIC26* mutations are also associated with mitochondrial myopathy, lactic acidosis (Beninca *et al*., 2021) as well as lethality and progeria-like phenotypes (Peifer-Weiß *et al*., 2023). We showed that cellular fatty acid synthesis, cholesterol biosynthesis and LD formation is promoted by MIC26 under high glucose conditions but that these pathways are conversely inhibited by MIC26 under normal glucose concentrations (**Fig 7H**). The important role of MIC26 in channelling nutrient excess from glucose into lipids underscores its reported links to obesity (Tian *et al*., 2017) and diabetes (Lamant *et al*., 2006) as it known that ectopic lipid accumulation is a common feature of the development of metabolic diseases including NAFLD and insulin resistance. Moreover, metabolism of glutamine via glutaminolysis is strongly impaired in the absence of MIC26. First, we discuss, how and why MIC26 promotes lipid anabolism in hyperglycemia and what is known from earlier studies in this context. Previously in mammalian cells, we characterised the role of MIC26, which contained a conserved apolipoprotein A1/A4/E family domain, in regulating mitochondrial ultrastructure and function (Koob *et al*., 2015). We showed that both an increase and decrease of MIC26 was detrimental to mitochondrial function indicating that optimal MIC26 amounts are essential for cellular homeostasis. Despite the demonstration of an increase of mitochondrial structural proteins, like MIC60, SAMM50 and MIC19, connected with upregulation of key metabolic pathways in mice fed with HFD compared to normal diet (Guo *et al*, 2013), the interplay of MICOS proteins, including MIC26, and metabolism is not clear. Since, classically apolipoproteins bind to lipids and mediate their transport in the bloodstream (Mehta & Shapiro, 2022), the presence of MIC26 in a non-classical environment like the IM raises various questions about its function. Interestingly, a previous report revealed a connection between increased levels of *MIC26* transcripts and nutrient conditions mimicked by oleic acid treatment (Wu *et al*, 2013). How does the loss of MIC26 alter central metabolic pathways including lipid metabolism in hyperglycemia? In this study, we found an increase of MIC26 in WT cells cultured in hyperglycemia. Concomitant to MIC26 increase, we found that MIC26 stimulates the formation of LDs when glucose is in excess. We demonstrate that MIC26 is essential for glucose utilisation and channelling glycolytic intermediates towards lipid anabolism regulating the accumulation of the LD content. This is supported by several findings including the determined levels of pyruvate and TCA cycle intermediates indicating that by boosting pyruvate levels, MIC26 further increases the amounts of the TCA cycle metabolites including citrate levels which serve as a precursor for cholesterol as well as FFA synthesis. This connection between lipid synthesis and MIC26 is further strengthened by earlier reports in the context of diabetes or obese models. Dogs fed with a HFD for 9 weeks (Philip-Couderc *et al*., 2003) and diabetic patients (Lamant *et al*., 2006) showed increased *Mic26* transcripts in the heart. Increased TAG and DAG were found upon *MIC26* overexpression in murine liver (Tian *et al*., 2017) and hearts (Turkieh *et al*., 2014) respectively showing modulatory roles of MIC26 in lipid metabolism. Our data reveals a major MIC26-dependent alteration of metabolite transporters of the mitochondrial IM and also metabolite levels. Thus, loss of MIC26 either alters the level, the activity, or the submitochondrial distribution of various metabolite transporters. In line with our data, the export of citrate from the mitochondrial matrix to the cytosol is presumably of particular importance. MIC26 could regulate metabolite exchange mechanistically either via protein-protein interactions of MIC26 to distinct metabolite transporters such as SLC25A12, by altering the accessibility of metabolites to various transporters due to altered cristae morphology. Overall, we propose that MIC26 regulates metabolite exchange between the cytosol and mitochondria and vice versa in a nutrient-dependent manner which is critical for adaptations to excess of glucose.

Under balanced nutrient conditions, MIC26 plays a different role compared to nutrient excess conditions. MIC26 decreases the key enzymes regulating the upper half of glycolytic pathway involved in ATP consumption phase. MIC26 prevents an increase of FFAs and G3P culminating in uncontrolled accumulation of LD content. In line with this, it was recently shown that the loss of MIC26 in BAT led to upregulation of glycolysis and fatty acid synthesis pathways (Guo *et al*., 2023). This was accompanied by impaired thermogenic activity of BAT, mitochondrial ultrastructure and function which reiterates the role of MIC26 in metabolic reprogramming. In normoglycemia, we found that the presence of MIC26 leads to a decrease of majority of the transcripts enzymes participating in cholesterol biosynthesis, including the sterol regulatory element binding transcription factor 2 (SREBP2) (**Fig S9C**) which is a master regulator of genes involved in sterol and fatty acid synthesis (Madison, 2016). However, we observed equal amounts of cholesterol in *MIC26* KO and WT cells under normoglycemia. Thus, MIC26 in normoglycemia facilitates equal metabolite distribution to either cholesterol biosynthesis or lipogenesis. This MIC26-mediated metabolic switch based on the amount/type of cellular fuel is essential for maintaining key metabolic pathways. A balanced amount of MIC26 is essential for how much glucose is channelled into lipid synthesis. In congruence with our observations, key lipid metabolism genes were altered upon *Mic26* overexpression with an interesting antagonistic regulation of *de novo* lipid synthesis genes depending on nutritional conditions (Tian *et al*., 2017). This study demonstrated an increase of important transcripts regulating lipid synthesis like ACACA, FASN and SCD in mice, overexpressing *Mic26*, fed with normal diet and a decrease when fed with HFD, compared to respective control mice. We further observed a decrease in CPT1A level and activity in *MIC26* KO cell lines as well as WT-H. CPT1A activity is known to be regulated either on a transcriptional level via peroxisome proliferator activated receptor α (PPARα) and peroxisome proliferator-activated receptor gamma coactivator 1 alpha (PGC-1α) or by allosteric inhibition through malonyl-CoA (López-Viñas *et al*, 2007; Song *et al*, 2010). A recent study demonstrated the downregulation of PPARα protein level in BAT in adipose tissue-specific *MIC26* KO mice (Guo *et al*., 2023). Further, we observed a high upregulation of ACACA enzyme, which converts acetyl-CoA to malonyl-CoA during *de novo* lipogenesis. Accordingly, it is possible that *MIC26* deletion in normoglycemia could on one hand reduce the expression of PPARα leading to decreased CPT1A expression and on the other hand increase malonyl-CoA formation leading to decreased CPT1A activity. Taken together, we used a multi-omics approach as well as a variety of functional assays to decipher that loss of MIC26 leads to an antagonistic regulation of glycolysis, lipid as well as cholesterol synthesis dependent on cellular nutritional stimulation.

Besides the antagonistic regulation mediated by MIC26 in different nutrient conditions, there are general roles of MIC26 in metabolic pathways which are independent of nutrient conditions. Among the MICOS proteins, proteins like MIC60 are considered as core components as *MIC60* deletion leads to a consistent loss of CJs (Kondadi *et al*., 2020a; Stephan *et al*., 2020) while the effect of *MIC26* deletion on loss of CJs varies with the cell line. Loss of CJs was observed in 143B (Koob *et al*., 2015) and HAP1 cells (Anand *et al*., 2020) in contrast to HeLa cells (Stephan *et al*., 2020). *MIC26* deletion in HepG2 cells in this study revealed a significant reduction of CJs when normalised to the cristae number, highlighting a possible major role of MIC26 in liver-derived cell lines. Concomitant to the reduction of CJs, we also observed alterations of vital transporters in the mitochondrial IM and OM. It was recently described that deletion of stomatin-like protein 2 (*SLP2*) leads to a drastic MIC26 degradation mediated by the YME1L protease (Naha *et al*, 2023). SLP2 was proposed as a membrane scaffold for PARL and YME1L named as the SPY complex (Wai *et al*, 2016). It is therefore conceivable that MIC26 could be concentrated in lipid-enriched nanodomains of the IM justifying its apolipoprotein nomenclature. When we checked the mitochondrial function upon *MIC26* deletion in HepG2 cells, we found that RCCs have enhanced respiratory capacity which was due to: a) increase in the levels of native RCCs as well as supercomplexes and b) increase in the activity of RCCs corresponding to increased RCC amounts. Thus, MIC26 could perform structural as well as functional roles which may or may not be mutually exclusive to the MICOS complex. A reduction of SLC25A12, an antiporter of cytoplasmic glutamate and mitochondrial aspartate, which is present in the IM was observed upon *MIC26* deletion independent of the nutritional status. We also found that SLC25A12 could be an interactor of MIC26 upon using standard interaction databases available online. Presumably, the interaction of SLC25A12 with MIC26 is important for the stability of the former. Such an intricate relationship between MIC26 and metabolite transporters in the IM makes it tempting to speculate that mitochondrial membrane remodelling is linked to its metabolic function. In fact, a closer look at the mitochondrial carrier family (SLC25) transcriptomics and proteomics data sets revealed that a majority of the SLC25 transporters were differentially regulated upon *MIC26* deletion in normoglycemia as well as hyperglycemia (**Fig S9A & B**). A prominent example is the reduction of SLC1A5 in *MIC26* KO when compared to WT in hyperglycemia as well as normoglycemia. A recent study showed that a variant of SLC1A5 present in the mitochondrial IM is responsible for transporting glutamine into the mitochondria (Yoo *et al*., 2020a). *MIC26* deletion leading to reduced total amounts of SLC1A5 also indicates reduced transport of glutamine into mitochondria. This was in line with an accumulation of glutamine upon loss of MIC26 at steady state. However, using glutamine tracing experiments we did not observe a decrease in labelled glutamate species but we observed an accumulation of α-KG. This could be due to the observed decrease in α-KG dehydrogenase resulting in reduced conversion of α-KG to other metabolites of the TCA cycle in particular under high glucose conditions. On the other hand, besides mitochondrial glutamine usage to fuel the TCA cycle, glutamine is known to be an essential source for nucleotide biosynthesis (Yoo *et al*, 2020b). *MIC26* KO cells showed a decreased growth rate (**Fig S9D-F**). Hence, glutamine accumulation and with that reduced conversion to nucleotides is a possible mechanism leading to growth deficiencies of *MIC26* KO cells. We found that the transcripts as well as protein levels of NUBPL were prominently downregulated upon *MIC26* deletion independent of the glucose concentrations of the cell culture media (**Fig S9G & H**). NUBPL was demonstrated to function as an assembly factor for complex I (Sheftel *et al*, 2009). Despite the prominent reduction of NUBPL, we did not find any discrepancy in complex I assembly or its activity most likely due to increased RCC amounts. We also found that the transcripts and protein levels of DHRS2 were significantly reduced in *MIC26* KO (**Fig S9I & J**). DHRS2 is implicated in reprogramming of lipid metabolism (Li *et al*, 2021) and was found to be downregulated in T2DM (De Silva K 2022).

We further found that hyperglycemia as well as *MIC26* deletion resulted in a fragmented mitochondrial morphology compared to WT-N. Mitochondrial dynamics and cellular metabolism including nutritional demands are closely interlinked (Mishra & Chan, 2016). Nutritional overload was associated with increased mitochondrial fragmentation (Yu *et al*, 2006) while starvation led to formation of a tubular mitochondrial network (Gomes *et al*, 2011). Further, mice lacking the ability to undergo mitochondrial fission by liver specific deletion of *Drp1* were protected from lipid accumulation in the liver as well as insulin resistance upon HFD feeding (Wang *et al*, 2015). Obesity is associated with increased mitochondrial fragmentation in multiple studies. A recent study showed that mitochondrial fragmentation is positively correlated to mitochondrial long chain FFA oxidation capacity via an increased activity of CPT1A (Ngo *et al*, 2023). A stronger membrane curvature resulting from mitochondrial fragmentation induces a conformational change leading to a decreased inhibitory binding ability of malonyl-CoA on CPT1 activity. Even though we observed mitochondrial fragmentation upon *MIC26* deletion, we did not observe increased FAO. This discrepancy could be explained by a reduction of CPT1A amount on one hand and likely increased production of malonyl-CoA on the other hand due to increased amounts of SLC25A1 and ACACA participating in fatty acid synthesis. Hence, deletion of MIC26, leading to mitochondrial fragmentation, contributes to ectopic cellular lipid accumulation but not FAO.

In sum, under balanced nutrient availability, we provide evidence that MIC26 is important to allow efficient metabolite channelling, mainly via glycolysis, thereby preventing unwanted channelling into lipogenesis. In addition, MIC26 is important to promote exactly the latter when glucose is in excess. This is important for cells to adapt to nutrient overload and explains earlier reports linking MIC26 to diabetes. We propose that MIC26 acts as a sensor and valve that opens towards lipid synthesis only when glucose is in excess. Future studies will have to decipher how changes in IM structure directly affect metabolite exchange and how this is regulated dynamically.

## Acknowledgments

A.K.K received Deutsche Forschungsgemeinschaft (DFG, German Research Foundation) Grant KO 6519/1-1. A.S.R received grants from DFG Graduiertenkolleg VIVID RTG 2576 and DFG SFB 1208, project B12 (ID 267205415). R.A received DFG Grant AN 1440/3-1. We thank Gisela Pansegrau and Andrea Borchardt for technical assistance in western blot and electron microscopy experiments, respectively. We are thankful for excellent technical support by Maria Graf, Elisabeth Klemp, and Katrin Weber (CMML). RNA-sequencing was performed at the Genomics & Transcriptomics facility, Biological and Medical Research Center (BMFZ), Medical Faculty, Heinrich-Heine-University Düsseldorf. Computational infrastructure and support were provided by the Centre for Information and Media Technology at Heinrich Heine University Düsseldorf. Proteome experiments were performed at the proteomics facility, BMFZ, HHU, Düsseldorf. Metabolite analyses were supported by the CEPLAS Plant Metabolism and Metabolomics Laboratory, which is funded by the DFG under Germanýs Excellence Strategy – EXC-2048/1 – project ID 390686111.

## Author Contributions

A.K.K. and A.S.R. conceptualized the research goals and the experiments of the study. M.L. planned, performed and analysed the results from majority of the experiments. R.N analysed and visualized proteomics and transcriptomics data. Y.S. planned, performed and analysed BN-PAGE and CN-PAGE. P.W. and A.P.M.W. performed and analysed metabolomics data. A.S. and K.S. performed and analysed proteomics data. P.P. and K.K. performed and analysed transcriptomics data. R.A. contributed with scientific and critical inputs to the study. A.K.K. and M.L. wrote the manuscript with input from all authors. A.K.K. supervised the study.

## Declaration of interests

The authors declare no competing interests

**S1 Figure. MIC26 loss leads to an opposing regulation of cholesterol biosynthesis pathway upon nutritional stimulation**

A. Overview of transcriptomics clustering analysis showing upregulated (red) and downregulated (blue) transcripts (without fold change or significance cut-offs). All sample replicates are represented (N = 4).

(B and C) Proteomics data represented by WikPathway enrichment using EnrichR analysis comparing pathways upregulated in normoglycemia (B) and downregulated in hyperglycemia (C) upon *MIC26* deletion compared to respective WT (N = 5). Arrows indicate increased levels of proteins participating in cholesterol synthesis and glycolysis pathways similar to those observed with transcriptomics data (Fig 2C and D). Differentially expressed proteins were considered statistically significant with a cut-off value of fold change of ±1.5 and Bonferroni correction *P* ≤ 0.05.

**S2 Figure. Loss of MIC26 leads to an opposing regulation of cholesterol biosynthesis pathway in normoglycemia and hyperglycemia**

(A and B) The transcripts of various enzymes regulating cholesterol synthesis are represented using Cytoscape software comparing log2FC data of *MIC26* KO and WT cell lines in normo-(A) and hyperglycemia (B) (N = 4). In *MIC26* KO cell lines, normoglycemia strongly increases transcripts of enzymes participating in cholesterol biosynthesis while an opposing effect is observed in hyperglycemia.

(C – N) Peptide abundances of various enzymes participating in cholesterol biosynthesis curated from proteomics data (N = 5).

(O) Metabolomics data reveals that cholesterol levels are exclusively decreased in *MIC26* KO at steady state in hyperglycemia compared to WT but not in normoglycemia (N = 3-4).

Data are represented as mean ± SEM (C-O). Statistical analysis was performed using one-way ANOVA with \**P* < 0.05, \*\**P* < 0.01, \*\*\**P* < 0.001, \*\*\*\**P* < 0.0001. N represents the number of biological replicates.

**S3 Figure. *MIC26* deletion causes opposing transcriptional regulation of genes involved in glycolysis**

(A and B) The transcripts of various enzymes participating in glycolysis are represented using Cytoscape software comparing log2FC data of *MIC26* KO and WT cell lines in normo- (A) and hyperglycemia (B) (N = 4).

(C and D) Glycolysis stress test Seahorse assay reveals a nonsignificant tendency towards increased glycolysis and glycolytic capacity upon *MIC26* KO in normoglycemic, but not in hyperglycemic conditions (N = 3).

Data are represented as mean ± SEM (C and D). Statistical analysis was performed using one-way ANOVA with \**P* < 0.05, \*\**P* < 0.01, \*\*\**P* < 0.001, \*\*\*\**P* < 0.0001. N represents the number of biological replicates.

**S4 Figure. Majority of free fatty acid species are antagonistically regulated upon *MIC26* deletion in normoglycemia and hyperglycemia when compared to respective to WT cells**

Detailed representation of abundances of various free fatty acid species in WT and *MIC26* KO cell lines cultured in normo- and hyperglycemia (N = 3-4).

Data are represented as mean ± SEM. Statistical analysis was performed using one-way ANOVA with \**P* < 0.05, \*\**P* < 0.01, \*\*\**P* < 0.001, \*\*\*\**P* < 0.0001. N represents the number of biological replicates.

**S5 Figure. *MIC26* deletion leads to alteration of key enzymes regulating lipid metabolism**

(A and B) The transcripts of mitochondrial long-chain fatty acid importer CPT1A are strongly decreased in WT-H and *MIC26* KO conditions, compared to WT-N, as shown from transcriptomics data (A) (N = 4) and quantitative PCR (B) (N = 3) analysis.

(C and D) Representative fatty acid oxidation assay analyzed using oxygen consumption rates of WT and *MIC26* KO HepG2 cells cultured in normoglycemia (C) and hyperglycemia (D) upon feeding either with BSA or palmitate (n = 8-12).

(E and F) Mitochondrial citrate malate exchanger (SLC25A1) is significantly increased upon *MIC26* deletion in normoglycemia compared to WT as detected using proteomics (E) (N = 5) and transcriptomics (F) (N = 4) data.

(G – M) Transcripts and available peptide abundances of key genes involved in lipid metabolism curated from transcriptomics (N = 4) and proteomics data (N = 5). Under normoglycemic conditions, loss of MIC26 increases the expression of ATP citrate lyase (G), acetyl-CoA carboxylase 1 (H and I), fatty acid synthase (J and K) and acetyl-CoA desaturase (L and M).

(N) Peptide abundances of glycerol kinase is increased in WT-H compared to WT-N but similar in *MIC26* KO-N and *MIC26* KO-H (N = 5).

Data are represented as mean ± SEM (A-N). Statistical analysis was performed using one-way ANOVA with \**P* < 0.05, \*\**P* < 0.01, \*\*\**P* < 0.001, \*\*\*\**P* < 0.0001. N represents the number of biological replicates and n the number of technical replicates.

**S6 Figure. TCA cycle enzymes are altered upon *MIC26* knockout**

(A – K) Representation of peptide abundances of various mitochondrial TCA cycle enzymes curated from proteomics data (N = 5).

(L – N) Peptide abundances of cytosolic enzymes involved in metabolite conversion (N = 5).

Data are represented as mean ± SEM (A-N). Statistical analysis was performed using one-way ANOVA with \**P* < 0.05, \*\**P* < 0.01, \*\*\**P* < 0.001, \*\*\*\**P* < 0.0001. N represents the number of biological replicates.

**S7 Figure. Mitochondrial glutamine and glutamate carriers are downregulated upon loss of MIC26**

(A) Transcripts of mitochondrial glutamate aspartate antiporter *SLC25A12* (A) are decreased upon *MIC26* deletion in normo- and hyperglycemia (N = 4).

(B and C) Transcripts (B) (N = 4) and peptide abundances (C) (N = 5) of cellular and mitochondrial glutamine importer SLC1A5 are significantly decreased upon *MIC26* deletion in both normo- and hyperglycemia in relation to respective WT cells.

(D) Peptide abundances of glutaminase (GLS) are unaltered upon loss of MIC26 compared to respective WT conditions.

(E) MIC26 interactome, based cumulatively on BioGRID, NeXtProt and IntAct databases, generated with Cytoscape software. From this study, downregulated transcripts comparing *MIC26* KO and WT in normoglycemic condition are highlighted in blue while upregulated transcripts are highlighted in red.

Data are represented as mean ± SEM (A-D). Statistical analysis was performed using one-way ANOVA with \**P* < 0.05, \*\**P* < 0.01, \*\*\**P* < 0.001, \*\*\*\**P* < 0.0001. N represents the number of biological replicates.

**S8 Figure. MIC26 maintains mitochondrial morphology and bioenergetics**

(A and B) Western Blots (A) and quantification (B) show a decrease of key mitochondrial fusion mediator MFN1 in *MIC26* KO cells while MFN2 was unchanged. Mitochondrial fission mediator DRP1 is decreased in *MIC26* KO-H compared to WT-H. OPA1 processing shows no significant changes upon *MIC26* deletion and nutritional status (N = 3-4).

(C and D) ATP production (C) and spare respiratory capacity (D) determined by mito stress test using Seahorse XF analyzer. Deletion of *MIC26* caused increased ATP production and decreased metabolic flexibility (indicated by SRC) in normoglycemia (N = 3).

Data are represented as mean ± SEM (B-D). Statistical analysis was performed using one-way ANOVA with \**P* < 0.05, \*\**P* < 0.01, \*\*\**P* < 0.001, \*\*\*\**P* < 0.0001. N represents the number of biological replicates.

**S9 Figure. *MIC26* deletion induces alterations of SLC25 mitochondrial carrier protein family expression and induces growth defects**

(A and B) Heat map overview from mean z-score of transcripts (A) (N = 4) and peptide abundances (B) (N = 5) of mitochondrial transporters belonging to SLC25 family.

(C) Transcript abundances of *SREBP2* (C) of WT and *MIC26* KO cells grown in normo- and hyperglycemia (N = 4).

(D – F) Proliferation of respective cell lines after 24 h (C), 48 h (D) and 72 h (E) determined using SRB assay normalized to WT-N (N = 4).

(G – J) Transcripts (N = 4) and peptide abundances (N = 5) of NUBPL (G and H) and DHRS2 (I and J) are respectively shown.

Data are represented as mean ± SEM (C-J). Statistical analysis was performed using one-way ANOVA with \**P* < 0.05, \*\**P* < 0.01, \*\*\**P* < 0.001, \*\*\*\**P* < 0.0001. N represents the number of biological replicates.

## Supplementary Tables

**Supplementary Table S1**

Raw data of targeted metabolomics at steady state including polar metabolites (sheet 1) and free fatty acids (FFAs) (sheet 2) indicating corresponding cell line, group, replicate, cell number, multiplication factor, measured compound name, internal standard (ISTD) name, measured total compound response, ISTD response, ratio of total compound response to ISTD response, relative response and technical procedure data. Relative response is calculated from compound response normalized to ISTD response and cell number multiplied by multiplication factor and used for data representation.

**Supplementary Table S2**

Raw data of targeted metabolomics from tracing experiments including AEC-MS (sheet 1) and GC-MS (sheet 2) data including cell line group analysed, compound isotopologue species, corrected ratio to naturally occurring isotopologues timepoint and technical information.

**Supplementary Table S3**

Proteomics raw data analysis including gene description, mean abundance ratio, adjusted *P*- value, mean abundances per group, total measured abundances of all replicate samples and technical data are represented in sheet 1. Filtered significantly (adj. *P*-value ≤ 0.05) altered peptide abundances with log2FC > ±1.5 for *MIC26* KO-N vs WT-N or *MIC26* KO-H and WT-H are represented in sheet 2 and 3 respectively. Detected peptides with less than three out of five hits in both of the compared groups were not considered.

**Supplementary Table S4**

Transcriptomics raw data analysis with sheet 1 representing raw data for all sample replicates, including gene description. Calculated log2FC, FC, *P*-Value, FDR adjusted *P*-Value and Bonferroni correction, as well as raw data from total reads, RPKM, TPM and CPM. DEGs filtered by Bonferroni correction ≤ 0.05 and log2FC > ±1.5 for *MIC26* KO-N vs WT-N and *MIC26* KO-H vs WT-H are represented in sheet 2 and 3 respectively. Sheet 4 is showing an overview of number of differentially expressed genes including respective cut-offs.

## Materials and Methods

### Key resources table

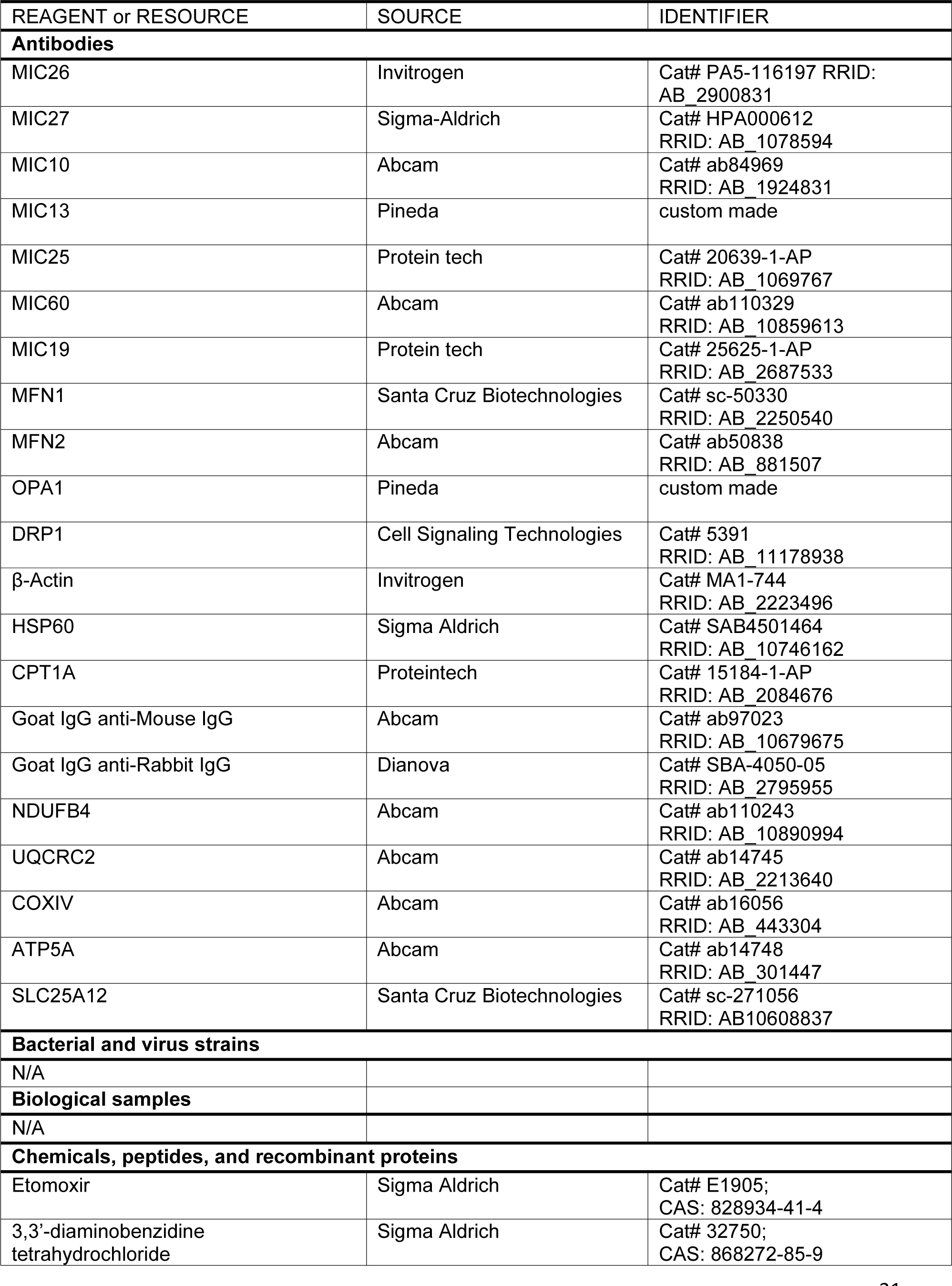
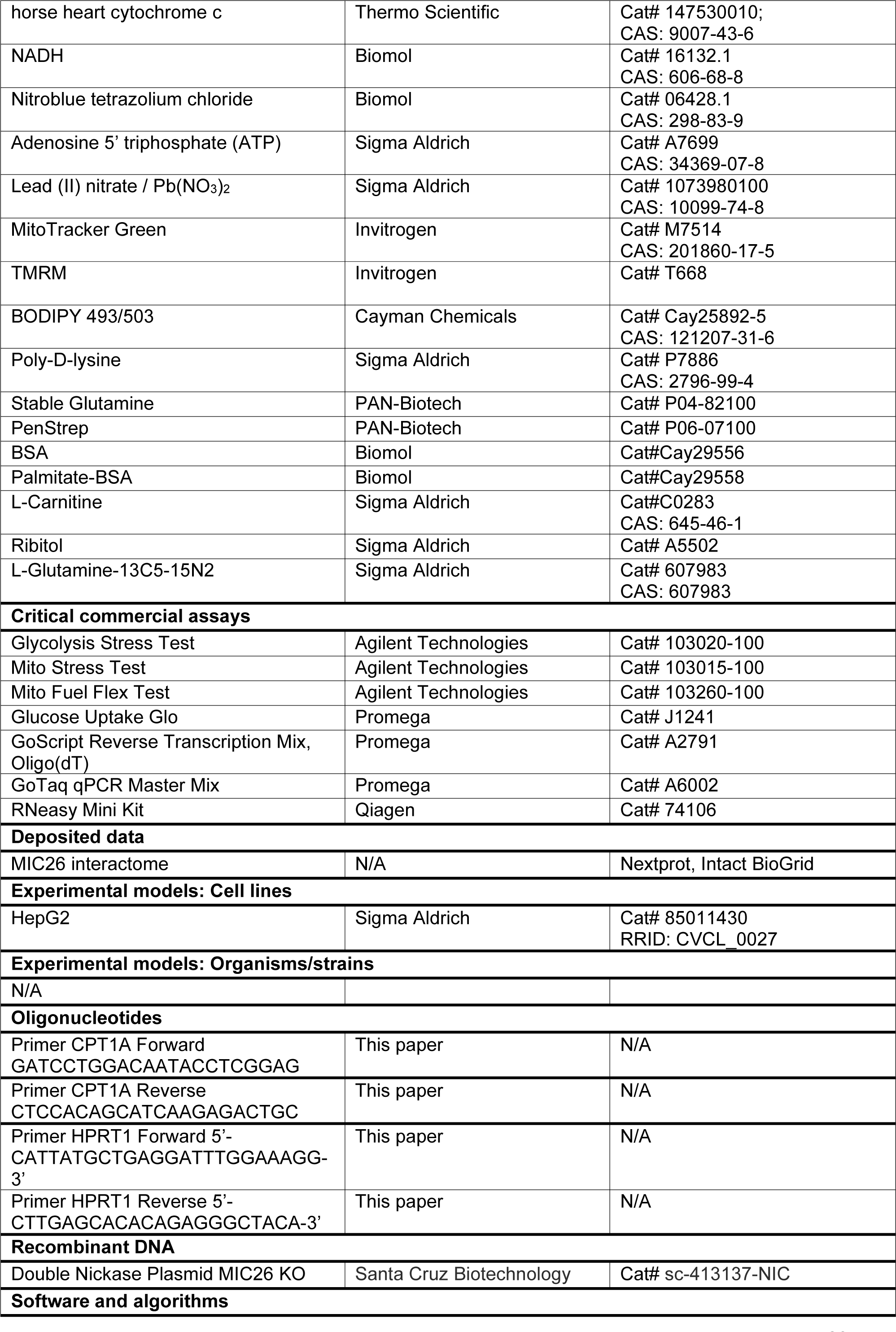
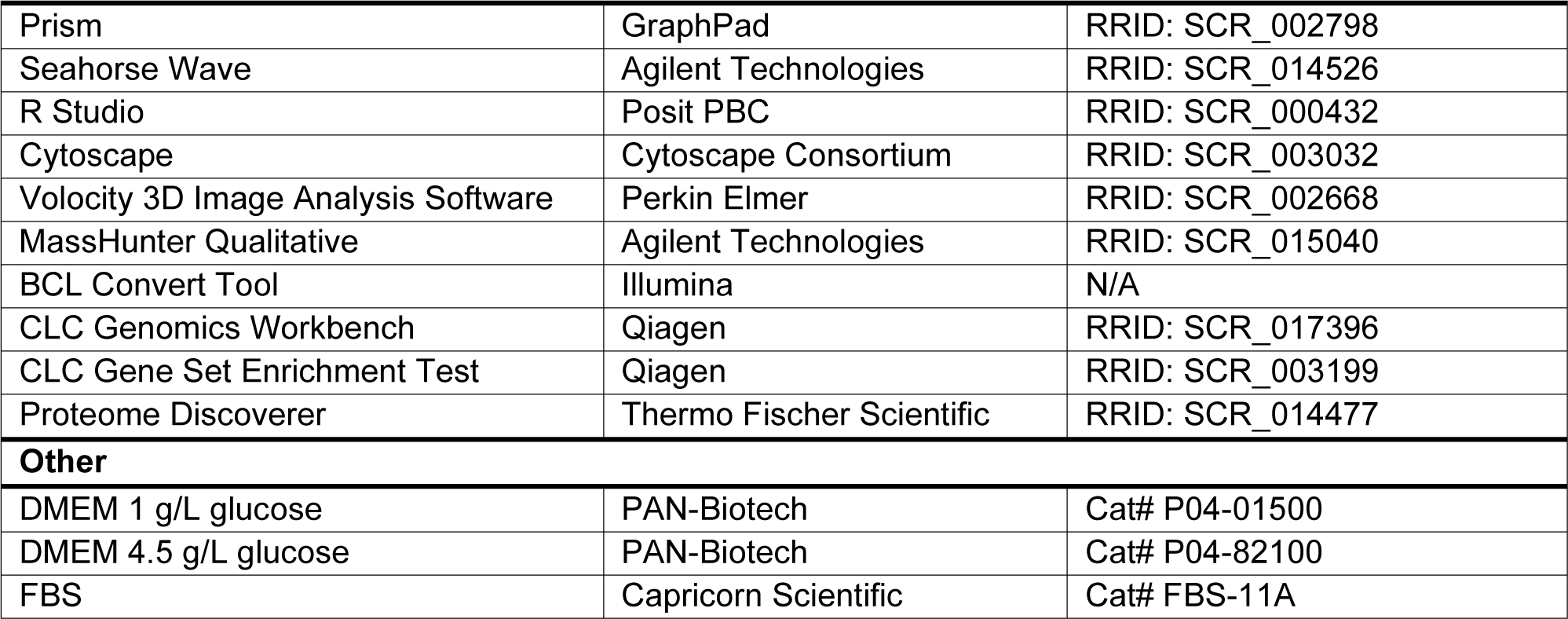

### Cell culture and treatment conditions

HepG2 cells were cultured in 1 g/L glucose DMEM (PAN-Biotech) supplemented with 10% FBS (Capricorn Scientific), 2 mM stable glutamine (PAN-Biotech) and penstrep (PAN-Biotech, penicillin 100 U/mL and 100 μg/mL streptomycin). Cells were grown at 37°C supplied with 5% CO_2_. *MIC26* HepG2 KO cells were generated using the double nickase method as described before (Lubeck *et al*., 2023). Cells cultured in standard growth media were divided equally into two cell culture flasks and grown in either 1 g/L glucose DMEM (normoglycemia) or 4.5 g/L glucose DMEM (hyperglycemia) (PAN-Biotech) supplemented with above-mentioned reagents. Cells were cultured in normoglycemia and hyperglycemia for a prolonged duration of three weeks. During the three weeks, cell splitting was carried out twice a week with the corresponding media.

### SDS gel electrophoresis and Western Blotting

After three washes with 2 mL DPBS (PAN-Biotech), the cells were harvested by scraping and resuspending in an appropriate volume of RIPA buffer (150 mM NaCl, 0.1 % SDS, 0.05 % Sodium deoxycholate, 1 % Triton-X-100, 1 mM EDTA, 1mM Tris, pH 7.4, 1x protease inhibitor (Sigma-Aldrich), PhosSTOP (Roche). Protein concentration was determined using DC^TM^ protein assay Kit (BIO-RAD, 5000116). SDS samples were prepared with Laemmli buffer and heated for 5 min at 95°C. Depending on the proteins investigated, a variety of SDS electrophoresis gels (8%, 10%, 12% or 15%) were used for running and separating protein samples. Subsequently, proteins were transferred onto nitrocellulose membranes and stained using Ponceau S (Sigma Aldrich). After destaining, nitrocellulose membranes were blocked with 5 % milk in 1x TBS-T for 1 h, washed three times with TBS-T and probed at 4°C overnight with the following primary antibodies: MIC26 (Invitrogen, 1:1000), MIC27 (Sigma-Aldrich, 1:2000), MIC10 (Abcam, 1:1000), MIC13 (Pineda custom-made, 1:1000), MIC25 (Proteintech, 1:1000), MIC60 (Abcam, 1:1000), MIC19 (Proteintech, 1:1000), MFN1 (Santa Cruz Biotechnologies, 1:1000), MFN2 (Abcam, 1:1000), OPA1 (Pineda custom-made, 1:1000), DRP1 (Cell Signaling Technology, 1:1000), β-Actin (Invitrogen, 1:2000), HSP60 (Sigma-Aldrich, 1:2000) and CPT1A (Proteintech, 1:1000). Goat IgG anti-Mouse IgG (Abcam, 1:10000) and Goat IgG anti-Rabbit IgG (Dianova, 1:10000) conjugated to HRP were used as secondary antibodies. The chemiluminescent signals were obtained using Signal Fire ECL reagent (Cell Signaling Technology) and VILBER LOURMAT Fusion SL equipment (Peqlab).

### Blue Native and Clear Native PAGE

5 x 10^6^ HepG2 cells were seeded onto 15 cm dishes and cell culture medium was replaced every two days until 80 % confluency was reached. Cells were washed three times with cold PBS, scraped and pelleted at 900 g, 4°C for 5 min. Cell pellets were resuspended in 1 mL lysis buffer (210 mM mannitol, 70 mM sucrose, 1 mM EDTA, 20 mM HEPES, 0,1 % BSA, 1x protease inhibitor) and incubated on ice for 10 min. Mitochondria were isolated by repetitive strokes of mechanical disruption using a 20G canula and sequential centrifugation steps at 1000 x g, 4°C for 10 min to remove cell debris and 10,000 x g, 4°C for 15 min to pellet mitochondria. Mitochondrial pellet was resuspended in BSA-free lysis buffer and protein concentration was determined using DC Protein Assay Kit.

For blue native page, 100 µg of mitochondria was solubilized for 1 h on ice using 2.5 g/g of digitonin to protein ratio. The samples were centrifuged for 20 min at 20,000 x g and 4°C to pellet insolubilized material. The supernatants were supplemented with loading buffer (50% glycerol, 8 g/g Coomassie to detergent ratio) and immediately loaded onto 3-13% gradient gel. Complexes were separated at 150 V, 15 mA for 16 h. Thereafter, protein complexes were transferred onto PVDF membrane and blocked overnight with 5 % milk in TBS-T at 4°C. For identification of relevant protein complexes, the membranes were decorated with the following antibodies: NDUFB4 (Abcam, 1:1000), UQCRC2 (Abcam, 1:1000), COXIV (Abcam, 1:1000) ATP5A (Abcam, 1:1000) Goat IgG anti-Mouse IgG (Abcam, 1:10000) and Goat IgG anti-Rabbit IgG (Dianova, 1:10000) conjugated to HRP. The chemiluminescent signals were obtained using Pierce™ SuperSignal™ West Pico PLUS chemiluminescent substrate reagent (Thermo Scientific) and VILBER LOURMAT Fusion SL equipment (Peqlab).

For clear native gels, 300 µg mitochondria was solubilized on ice for 1 h with 2.5 g/g digitonin to protein ratio. The samples were centrifuged for 20 min at 20,000 x g and 4°C to pellet insolubilized material. The supernatants were supplemented with loading buffer (50% glycerol, 0.01 % Ponceau S) and immediately loaded onto 3-13% gradient gels. Complexes were separated at 150 V, 15 mA for 16 h. To assess complex in-gel activity, the gel slices were incubated in respective buffer solutions for several hours at room temperature. For complex I activity, the gel was incubated in 5 mM Tris-HCl (pH 7.4), 0.1 mg/mL NADH and 2.5 mg/mL nitro blue tetrazolium chloride (NBT). For complex III, the gel was incubated in 50 mM sodium phosphate buffer (pH 7.2), 0.1 % 3,3’-diaminobenzidine tetrahydrochloride (DAB). To assess complex IV activity, the gel was incubated in 50 mM sodium phosphate buffer (pH 7.2), 0.05 % DAB and 50 µM horse heart cytochrome *c* and for complex V, the gel was incubated in 35 mM Tris-base, 270 mM glycine, 14 mM MgSO_4_, 0.2 % (w/v) Pb(NO_3_)_2_ and 8 mM ATP.

### RNA isolation and quantification

Total RNA was extracted from cell pellets using RNeasy Mini Kit (Qiagen) according to the manufacturer’s protocol. RNA quality and quantity were assessed using BioSpectrometer (Eppendorf). cDNA synthesis from 5 µg RNA was performed using the GoScript^TM^ Reverse Transcriptase Kit (Promega). Next, quantitative real-time PCR was performed in Rotor Gene 6000 (Corbett Research) using GoTagR qPCR Master Mix (Promega) according to manufacturer’s instructions with the following primers:

1. *CPT1A*: Forward: 5’- GATCCTGGACAATACCTCGGAGC-3’ Reverse: 5’- CTCCACAGCATCAAGAGACTGC-3’
2. *HPRT1* (Housekeeping gene): Forward: 5’-CATTATGCTGAGGATTTGGAAAGG-3’ Reverse: 5’-CTTGAGCACACAGAGGGCTACA-3’ C_t_ values were normalized to housekeeping gene *HPRT1* followed by normalization of ΔC_t_ values to average ΔC_t_ of WT-N control group.

### Transcriptomics

Cells were seeded in quadruplicates onto 10 cm dishes in corresponding cell culture media and medium was replaced every two days until 80 % cell confluency was obtained. For preparation of RNA, cells were washed three times with cold PBS and subsequently scraped and pelleted. RNA isolation from cell pellets was performed using RNeasy Mini Kit (Qiagen) including DNase digestion according to the manufacturer’s protocol. Sample concentration was determined and 1 µg RNA was aliquoted for transcriptomics analysis. Total RNA samples were quantified (Qubit RNA HS Assay, Thermo Fisher Scientific, MA, USA) and quality measured by capillary electrophoresis using the Fragment Analyzer and the ’Total RNA Standard Sensitivity Assay’ (Agilent Technologies, Inc. Santa Clara, CA, USA). All samples in this study showed RNA Quality Numbers (RQN) with a mean of 10.0. The library preparation was performed according to the manufacturer’s protocol using the ’VAHTS™ Stranded mRNA- Seq Library Prep Kit’ for Illumina®. Briefly, 700 ng total RNA were used as input for mRNA capturing, fragmentation, the synthesis of cDNA, adapter ligation and library amplification. Bead purified libraries were normalized and finally sequenced on the NextSeq2000 system (Illumina Inc. San Diego, CA, USA) with a read setup of 1×100 bp. The BCL Convert Tool (version 3.8.4) was used to convert the bcl files to fastq files as well for adapter trimming and demultiplexing.

Data analyses on fastq files were conducted with CLC Genomics Workbench (version 22.0.2, Qiagen, Venlo, Netherlands). The reads of all probes were adapter trimmed (Illumina TruSeq) and quality trimmed (using the default parameters: bases below Q13 were trimmed from the end of the reads, ambiguous nucleotides maximal 2). Mapping was done against the *Homo sapiens* (hg38; GRCh38.107) (July 20, 2022) genome sequence. After grouping of samples (four biological replicates) according to their respective experimental conditions, the statistical differential expression was determined using the CLC differential expression for RNA-Seq tool (version 2.6, Qiagen, Venlo, Netherlands). The resulting *P* values were corrected for multiple testing by FDR and Bonferroni-correction. A *P* value of ≤0.05 was considered significant. The CLC gene set enrichment test (version 1.2, Qiagen, Venlo, Netherlands) was done with default parameters and based on the GO term ’biological process’ (*H. sapiens*; May 01, 2021).

The data discussed in this publication have been deposited in NCBI’s Gene Expression Omnibus (Edgar *et al*, 2002) and are accessible through GEO Series accession number GSE248848.

### Proteomics

Cells were seeded in quintuplicates onto 10 cm dishes in corresponding cell culture media and medium was replaced every two days until 80 % cell confluency was obtained. Cells were washed four times with PBS, scraped and pelleted in a pre-weighed Eppendorf tube. After complete removal of PBS, cells were immediately frozen in liquid nitrogen and sample weight was determined for normalization. Proteins were extracted from frozen cell pellets as described elsewhere (Poschmann *et al*, 2014). Briefly, cells were lysed and homogenized in urea buffer with a TissueLyser (Qiagen) and supernatants were collected after centrifugation for 15 min at 14,000 x g and 4°C. Protein concentration was determined by means of Pierce 660 nm protein assay (Thermo Fischer Scientific). For LC-MS analysis, a modified magnetic bead-based sample preparation protocol according to Hughes and colleagues was applied (Hughes *et al*, 2019). Briefly, 20 µg total protein per sample was reduced by adding 10 µL 100 mM DTT (dithiothreitol) and shaking for 20 min at 56°C and 1000 rpm, followed by alkylation with the addition of 13 µL 300 mM IAA and incubation for 15 min in the dark. A 20 µg/µL bead stock of 1:1 Sera-Mag SpeedBeads was freshly prepared and 10 µL was added to each sample. Afterwards, 84 µL ethanol was added and incubated for 15 min at 24°C. After three rinsing steps with 80% EtOH and one rinsing step with 100% ACN, beads were resuspended in 50 mM TEAB buffer and digested with final 1:50 trypsin at 37°C and 1,000 rpm overnight. Extra- digestion was carried out by adding trypsin (final 1:50) and shaking at 37°C and 1000 rpm for 4 h. The supernatants were collected and 500 ng of each sample digest was subjected to LC- MS.

For the LC-MS acquisition, an Orbitrap Fusion Lumos Tribrid Mass Spectrometer (Thermo Fisher Scientific) coupled to an Ultimate 3000 Rapid Separation liquid chromatography system (Thermo Fisher Scientific) equipped with an Acclaim PepMap 100 C18 column (75 µm inner diameter, 25 cm length, 2 µm particle size from Thermo Fisher Scientific) as separation column and an Acclaim PepMap 100 C18 column (75 µm inner diameter, 2 cm length, 3 µm particle size from Thermo Fisher Scientific) as trap column was used. A LC-gradient of 180 min was applied. Survey scans were carried out over a mass range from 200-2,000 m/z at a resolution of 120,000. The target value for the automatic gain control was 250,000 and the maximum fill time 60 ms. Within a cycle time of 2 s, the most intense peptide ions (excluding singly charged ions) were selected for fragmentation. Peptide fragments were analysed in the ion trap using a maximal fill time of 50 ms and automatic gain control target value of 10,000 operating in rapid mode. Already fragmented ions were excluded for fragmentation for 60 seconds.

Data analysis was performed with Proteome Discoverer (version 2.4.1.15, Thermo Fisher Scientific). All RAW files were searched against the human Swissprot database (Download: 23.01.2020) and the Maxquant Contaminant database (Download: 20.02.2021), applying a precursor mass tolerance of 10 ppm and a mass tolerance of 0.6 Da for fragment spectra. Methionine oxidation, N-terminal acetylation, N-terminal methionine loss and N-terminal methionine loss combined with acetylation were considered as variable modifications, carbamidomethylation as static modification as well as tryptic cleavage specificity with a maximum of two missed cleavage sites. Label-free quantification was performed using standard parameters within the predefined workflow. Post processing, proteins were filtered to 1% FDR and a minimum of 2 identified peptides per protein. The mass spectrometry proteomics data have been deposited to the ProteomeXchange Consortium via the PRIDE (Perez-Riverol *et al*, 2022) partner repository with the dataset identifier PXD047246.

### Metabolomics

Metabolites were analyzed by gas chromatography (GC) and anion exchange chromatography (AEC) coupled to mass spectrometry (MS). 1.5 x 10^6^ cells were seeded in quadruplicates onto 6 cm dishes and cultured in the corresponding media overnight. For glutamine tracing experiments, medium was replaced with corresponding growth media containing 2 mM labeled glutamine [U-^13^C_5_, ^15^N_2_] (Sigma-Aldrich) either for 30 min or 6 h prior to cell harvesting. For metabolite extraction, cells were washed five times with ice-cold isotonic NaCl solution (0.9 %), followed by scraping of cells in 1 mL ice-cold MeOH. Cells were transferred to a 15 mL tube and diluted with 1 mL MilliQ water. Cell suspension was immediately frozen in liquid nitrogen.

After thawing on ice, 0.5 mL MilliQ water was added supplemented with 10 µM internal standard ribitol (Sigma Aldrich) for polar metabolite analysis. After that 1.5 mL MTBE was added containing 5.4 μL heptadecanoic acid (1mg/ml) as internal standard for free fatty acid analysis. After repetitive mixing, samples were incubated on ice for 10 min. Subsequently, polar and nonpolar phases were separated by centrifugation at 4000 x g for 10 min at 4°C. The apolar phase was collected, frozen at -80°C and used for free fatty acid analysis. The aqueous phase was diluted with MilliQ water to decrease the organic proportion below 15 %. The sample was then frozen at -80°C, dried by lyophilization reconstituted in 500 µL MilliQ water and filtered prior to analysis.

For GC-MS, 100 µL was dried by vacuum filtration. Metabolite analysis was conducted using a 7890B gas chromatography system connected to a 7200 QTOF mass spectrometer (Agilent Technologies) as described previously (Shim *et al*, 2019). In brief, methoxyamine hydrochloride and N-methyl-N-(trimethylsilyl)trifluoroacetamide were subsequently added to the dried sample to derivatize functional groups of polar compounds. With an injection volume of 1 µL, samples were introduced into the GC-MS system and compounds were separated on a HP-5MS column (30m length, 0.25mm internal diameter and 0.25µm film thickness). The software MassHunter Qualitative (v b08, Agilent Technologies) was used for compound identification by comparing mass spectra to an in-house library of authentic standards and to the NIST14 Mass Spectral Library (https://www.nist.gov/srd/nist-standard-reference-database-1a-v14). Peak areas were integrated using MassHunter Quantitative (v b08, Agilent Technologies) and normalized to the internal standard ribitol and cell number. To determine the ^13^C and ^15^N incorporation, isotopologues for individual fragments were analyzed according to the number of possible incorporation sites. The normalized peak areas were corrected for the natural abundance using the R package IsoCorrectoR (Heinrich *et al*, 2018).

For the analysis of anionic compounds by AEC-MS, samples were diluted with MilliQ water (1:2 v/v). Measurements were performed using combination of a Dionex ICS-6000 HPIC and a high field Thermo Scientific Q Exactive Plus quadrupole-Orbitrap mass spectrometer (both Thermo Fisher Scientific) as described earlier with minor modifications (Curien *et al*, 2021). 10 µL of sample was injected via a Dionex AS-AP autosampler in push partial mode. Anion exchange chromatography was conducted on a Dionex IonPac AS11-HC column (2 mm X 250 mm, 4 µm particle size, Thermo Scientific) equipped with a Dionex IonPac AG11-HC guard column (2 mm X 50 mm, 4 µm, Thermo Scientific) at 30°C. The mobile phase was established using an eluent generator with a potassium hydroxide cartridge to produce a potassium hydroxide gradient. The column flow rate was set to 380 μL min^-1^ with a starting KOH concentration of 5 mM for one minute. The concentration was increased to 85 mM within 35 min and held for 5 min. The concentration was immediately reduced to 5 mM and the system equilibrated for 10 min. Spray stability was achieved with a makeup consisting of methanol _-1_ with 10 mM acetic acid delivered with 150 μL min^-1^ by an AXP Pump. The electro spray was achieved in the ESI source using the following parameters: sheath gas 30, auxiliary gas 15, sweep gas 0, spray voltage - 2.8 kV, capillary temperature 300°C, S-Lens RF level 45, and auxiliary gas heater 380°C. For the untargeted approach, the mass spectrometer operated in a combination of full mass scan and a data-dependent Top5 MS2 (ddMS2) experiment. The full scan (60-800 m/z) was conducted with a resolution of 140,000 and an automatic gain _6_ control (AGC) target of 10 ions with a maximum injection time of 500 ms. The Top5 ddMS2 _5_ experiment was carried out with a resolution of 17,500 and an AGC target of 10 and a maximum IT of 50 ms. The stepped collision energy was used with the steps (15, 25, 35) to create an average of NCE 25. Data analysis was conducted using Compound Discoverer (version 3.1, Thermo Scientific) using the “untargeted Metabolomics workflow” for steady state analysis. Compound identification was achieved on the level of mass accuracy (MS1 level), fragment mass spectra matching (MS2 level) and retention time comparison with authentic standards. For the enrichment analysis with stable heavy isotopes, the standard workflow for “stable isotope labelling” was chosen with the default settings 5 ppm mass tolerance, 30 % intensity tolerance and 0.1 % intensity threshold for isotope pattern matching and a maximum exchange rate was of 95%.

For free fatty acid analysis via GC-MS, lipids were hydrolysed and free fatty acids were methylated to fatty acid methyl esters (FAMEs). To do so, the organic phase was transferred into a glass vial and dried under a stream of nitrogen gas. The dried sample was resuspended in 1 mL of methanolic hydrochloride (MeOH/3 N HCl) and incubated at 90°C for 1 h. One mL of hexane and 1 mL of NaCl solution (1%) were added before centrifugation at 2000 *g* for 5 min. The FAME-containing organic phase (top layer) was collected in a clean glass vial and stored at −20°C until measurement as described recently (Vasilopoulos *et al*, 2023).

### Quantification of mitochondrial morphology, membrane potential (ΔΨ_m_) and cellular lipid droplets

HepG2 cells (0.25 x 10^6^ cells) were seeded onto 35 mm Poly-D-Lysine-coated (50 µg/ml) live-imaging dishes (MATTEK P35G-1.5-14-C) and incubated for 24 h at 37°C, 5 % CO_2_ in the corresponding normoglycemic or hyperglycemic media. The assessment of mitochondrial morphology, ΔΨ_m_ and cellular lipid droplets was performed by addition of MitoTracker Green (Invitrogen, 200 nM), TMRM (Invitrogen, 50 nM), BODIPY 493/503 (Cayman Chemicals, 10 µM) respectively for 30 min at 37°C, followed by washing thrice. Live-cell microscopy was performed using a spinning disc confocal microscope (PerkinElmer) equipped with a 60x oil-immersion objective (N.A = 1.49) and a Hamamatsu C9100 camera (1000 X 1000 pixel). The cells were maintained at 37°C in DMEM supplemented with 10 mM HEPES for the imaging duration. MitoTracker Green and BODIPY 493/503 were excited with a 488 nm laser while TMRM was excited with a 561 nm laser. The images were obtained at emission wavelength of 527 nm (W55) and 615 nm (W70) for 488 nm and 561 nm excitation respectively. The cell population was classified into tubular, intermediate and fragmented mitochondrial morphology based on the majority of mitochondria belonging to the respective class. Cells classified as tubular and fragmented contained mostly long tubular and short fragments respectively whereas cells classified as intermediate had a mixture of mostly short pieces, few long tubes as well as fragmented mitochondria. Volocity image analysis software was used for the quantification regarding ΔΨ_m_ and lipid droplets. The total fluorescence intensities of TMRM and BODIPY were obtained per cell after respective background subtraction. Each cell was manually demarcated by drawing a ROI. Lipid droplet number within a ROI was obtained automatically using find spots by setting threshold of brightest spot within a radius of 0.5 µm and compartmentalization to ROI.

### Glucose Uptake Assay

3×10^4^ HepG2 cells were seeded in triplicates onto a dark 96-well plate overnight and in parallel onto a clear-96 well plate for cell normalization. Cellular glucose uptake was measured using Glucose Uptake-Glo^TM^ Assay kit (Promega), according to the manufacturer’s protocol. Luminescence was measured by microplate reader (CLARIOstar Plus, BMG LABTECH) with 1 s integration after 1 h of incubation. Normalization was performed using Hoechst staining and mean of signal intensity was used for normalizing luminescence intensities. Luciferase signals were normalized to WT-N measurement.

### Mitochondrial respirometry

A variety of respirometry experiments were performed using Seahorse XFe96 Analyzer (Agilent). HepG2 cells were seeded onto Poly-D-Lysine-coated (50 µg/ml) Seahorse XF96 cell culture plate (Agilent) at a density of 3.0 x 10^4^ cells per well. For mitochondrial stress test, mitochondrial fuel flexibility test and glycolysis stress test, cells were incubated overnight in standard growth media. For fatty acid oxidation (FAO) test, standard growth medium was replaced by serum-deprived growth medium (DMEM without glucose, pyruvate and glutamine), containing 1 % FBS, 0.5 mM glucose, 0.5 mM L-Carnitine (Sigma Aldrich) and 1.0 mM glutamine 10 h after cell seeding and incubated overnight.

Prior to performing the assay, old medium was removed and cells were washed twice after which cells were supplemented with the corresponding assay media followed by 45 min CO_2_- free incubation. Mitochondrial stress test was performed using Seahorse assay media (Agilent) supplemented with 10 mM glucose, 2 mM stable glutamine and 1 mM sodium pyruvate.

Mitochondrial oxygen consumption was measured after sequential addition of oligomycin (1 µM), FCCP (0.25 µM) and rotenone/antimycin (0.5 µM) according to the manufacturer’s protocol. Mitochondrial fuel flexibility test was performed using Seahorse assay media containing 10 mM glucose, 2 mM stable glutamine and 1 mM sodium pyruvate. After initial acquisition of basal respiration, glucose, glutamine and FAO dependency and capacity was assessed according to manufacturer’s protocol by sequential incubation with UK5099 (2 µM) and Etomoxir (4 µM) / BPTES (3 µM), BPTES (3 µM) and Etomoxir (4 µM) / UK5099 (2 µM) or Etomoxir (4 µM) and UK5099 (2 µM) / BPTES (3 µM) respectively. Glycolysis stress test was performed in Seahorse assay media supplemented with 2 mM glutamine. After 15 min of basal ECAR determination, glycolysis was induced by addition of glucose (10 mM), followed by oligomycin (1 µM) and lastly 2-DG (50 mM). For assessment of FAO, cells were pretreated with Seahorse assay media containing BSA (Biomol, 200 µM) or Palmitate (Biomol, 200 µM). FAO was measured by sequential addition of etomoxir (Sigma Aldrich, 4 µM) or media and mitochondrial stress test kit chemicals oligomycin (1.5 µM), FCCP (1 µM), rotenone/antimycin A (0.5 µM). Cell numbers were normalized using Hoechst (10 ug/mL) staining intensity assessed by microplate reader (Tecan M200 pro). Data were analyzed using wave software (Agilent) and Microsoft Excel.

### Electron Microscopy

4 x 10^6^ HepG2 cells were grown overnight in 10 cm petri dishes at 37°C with 5% CO_2_ in the corresponding treatment media. Cells were fixed using 3 % glutaraldehyde, 0.1 M sodium cacodylate buffer at pH 7.2 and subsequently pelleted. Cell pellets were washed in fresh 0.1 M sodium cacodylate buffer at pH 7.2 and embedded in 3 % low melting agarose. Cells were stained using 1% osmium tetroxide for 50 min, washed twice with 0.1 M sodium cacodylate buffer and once using 70% ethanol for 10 min each. Thereafter, cells were stained using 1% uranyl acetate/1% phosphotungstic acid in 70% ethanol for 1 h. Stained samples were embedded in spur epoxy resin for polymerization at 70°C for 24 hours. Ultrathin sections were prepared using a microtome and imaged on a transmission electron microscope (Hitachi, H600) at 75 V equipped with Bioscan 792 camera (Gatan). Image analysis was performed using ImageJ software.

### Sulforhodamine B (SRB) assay

Cell viability was assessed by SRB colorimetry assay. 2.5 x 10^4^ HepG2 cells were seeded in 24 well plates and incubated for 24 h, 48 h or 72 h. Subsequently, cells were washed with PBS and fixed with 10% (w/v) cold trichloroacetic acid solution (500 μL/well) for 1 h at 4°C. After washing five times with MilliQ water, cells were dried at RT overnight. Fixed cells were stained with SRB solution (0.4% (w/v) in 1% acetic acid, 300 μl/well) for 15 min at RT, washed five times with 1% acetic acid and dried at RT for 1 h. SRB extraction was performed by addition of 400 μL TRIS-Base (10 mmol/l) per well. The absorbance was measured, after 5 min of shaking, at 492 nm and 620 nm using a microplate reader (Tecan M200 pro). Total intensity was calculated from signal intensity at 492 nm after background subtraction of 620 nm intensity. Proliferation was normalized to WT-N.

### Statistics and data representation

Data are represented as mean ± standard error mean (SEM). Statistical significance was determined by one-way ANOVA followed by Šídák’s test for multiple comparisons of selected pairs with \**P*-value ≤ 0.05, \*\**P*-value ≤ 0.01, \*\*\**P*-value ≤ 0.001, \*\*\*\**P*-value ≤ 0.0001. Data analysis was performed using Microsoft Excel. Data representation and statistical analysis was performed using GraphPad Prism.

## Notes

### Competing Interest Statement

The authors have declared no competing interest.

