## Supplementary Figure 1 for "Mitochondrial Apolipoprotein MIC26 is a metabolic rheostat regulating central cellular fuel pathways"

**S1 Figure**

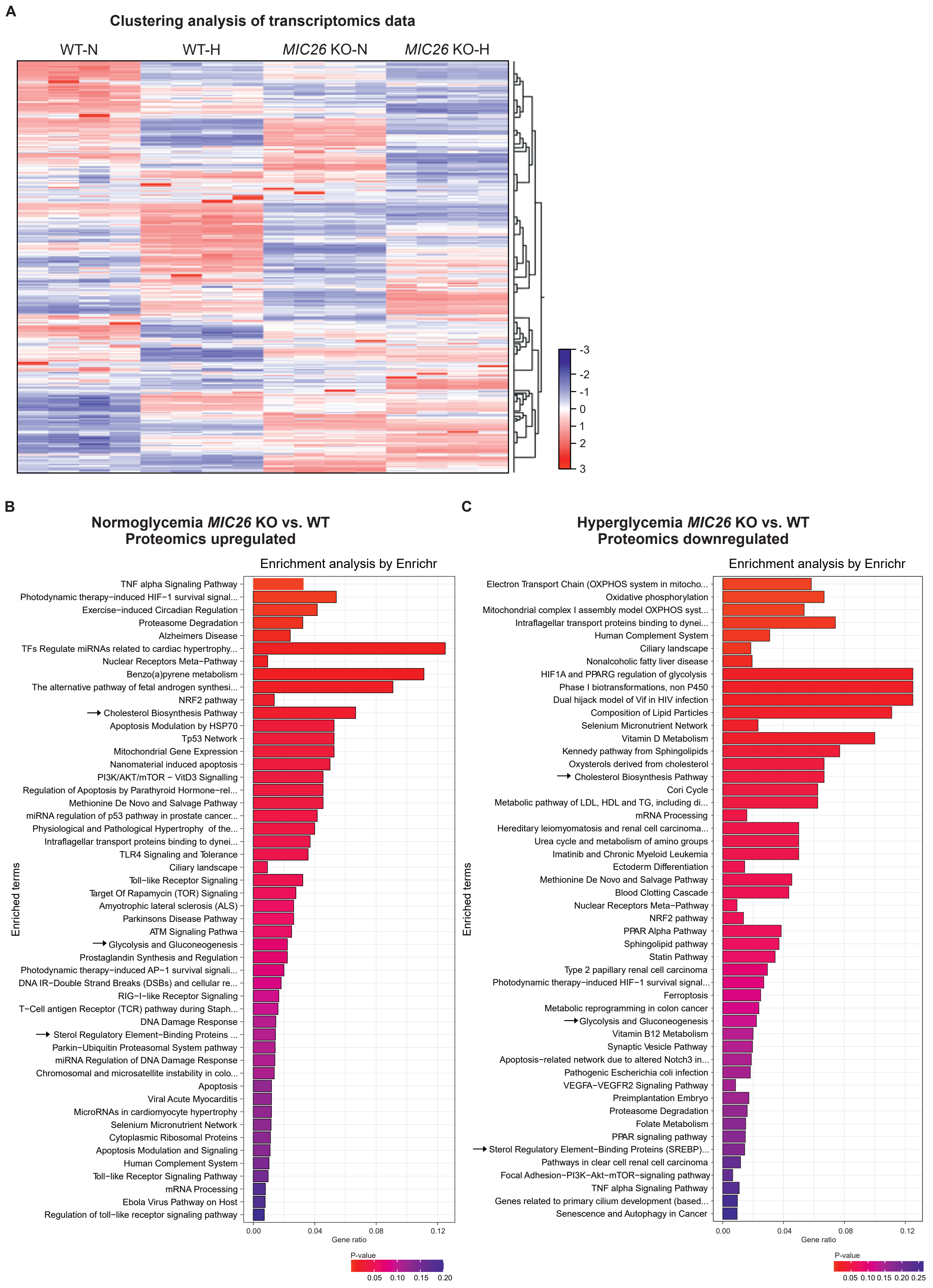
A
Clustering analysis of transcriptomics data
WT-N
WT-H
*MIC26* KO-N
*MIC26* KO-H
-3
-2
-1
0
1
2
3
B
Normoglycemia *MIC26* KO vs. WT
Proteomics upregulated
Enrichment analysis by Enrichr
TNF alpha Signaling Pathway
Photodynamic therapy−induced HIF−1 survival signal...
Exercise−induced Circadian Regulation
Proteasome Degradation
Alzheimers Disease
TFs Regulate miRNAs related to cardiac hypertrophy...
Nuclear Receptors Meta−Pathway
Benzo(a)pyrene metabolism
The alternative pathway of fetal androgen synthesi...
NRF2 pathway
Cholesterol Biosynthesis Pathway
Apoptosis Modulation by HSP70
Tp53 Network
Mitochondrial Gene Expression
Nanomaterial induced apoptosis
PI3K/AKT/mTOR − VitD3 Signalling
Regulation of Apoptosis by Parathyroid Hormone−rel...
Methionine De Novo and Salvage Pathway
miRNA regulation of p53 pathway in prostate cancer...
Physiological and Pathological Hypertrophy of the...
Intraflagellar transport proteins binding to dynei...
TLR4 Signaling and Tolerance
Ciliary landscape
Toll−like Receptor Signaling
Target Of Rapamycin (TOR) Signaling
Amyotrophic lateral sclerosis (ALS)
Parkinsons Disease Pathway
ATM Signaling Pathwa
Glycolysis and Gluconeogenesis
Prostaglandin Synthesis and Regulation
Photodynamic therapy−induced AP−1 survival signali...
DNA IR−Double Strand Breaks (DSBs) and cellular re...
RIG−I−like Receptor Signaling
T−Cell antigen Receptor (TCR) pathway during Staph...
DNA Damage Response
Sterol Regulatory Element−Binding Proteins ...
Parkin−Ubiquitin Proteasomal System pathway
miRNA Regulation of DNA Damage Response
Chromosomal and microsatellite instability in colo...
Apoptosis
Viral Acute Myocarditis
MicroRNAs in cardiomyocyte hypertrophy
Selenium Micronutrient Network
Cytoplasmic Ribosomal Proteins
Apoptosis Modulation and Signaling
Human Complement System
Toll−like Receptor Signaling Pathway
mRNA Processing
Ebola Virus Pathway on Host
Regulation of toll−like receptor signaling pathway
Enriched terms
0.00
0.04
0.08
0.12
Gene ratio
P-value
0.05
0.10
0.15
0.20
C
Hyperglycemia *MIC26* KO vs. WT
Proteomics downregulated
Enrichment analysis by Enrichr
Electron Transport Chain (OXPHOS system in mitocho...
Oxidative phosphorylation
Mitochondrial complex I assembly model OXPHOS syst...
Intraflagellar transport proteins binding to dynei...
Human Complement System
Ciliary landscape
Nonalcoholic fatty liver disease
HIF1A and PPARG regulation of glycolysis
Phase I biotransformations, non P450
Dual hijack model of Vif in HIV infection
Composition of Lipid Particles
Selenium Micronutrient Network
Vitamin D Metabolism
Kennedy pathway from Sphingolipids
Oxysterols derived from cholesterol
Cholesterol Biosynthesis Pathway
Cori Cycle
Metabolic pathway of LDL, HDL and TG, including di...
mRNA Processing
Hereditary leiomyomatosis and renal cell carcinoma...
Urea cycle and metabolism of amino groups
Imatinib and Chronic Myeloid Leukemia
Ectoderm Differentiation
Methionine De Novo and Salvage Pathway
Blood Clotting Cascade
Nuclear Receptors Meta−Pathway
NRF2 pathway
PPAR Alpha Pathway
Sphingolipid pathway
Statin Pathway
Type 2 papillary renal cell carcinoma
Photodynamic therapy−induced HIF−1 survival signal...
Ferroptosis
Metabolic reprogramming in colon cancer
Glycolysis and Gluconeogenesis
Vitamin B12 Metabolism
Synaptic Vesicle Pathway
Apoptosis−related network due to altered Notch3 in...
Pathogenic Escherichia coli infection
VEGFA−VEGFR2 Signaling Pathway
Preimplantation Embryo
Proteasome Degradation
Folate Metabolism
PPAR signaling pathway
Sterol Regulatory Element−Binding Proteins (SREBP)...
Pathways in clear cell renal cell carcinoma
Focal Adhesion−PI3K−Akt−mTOR−signaling pathway
TNF alpha Signaling Pathway
Genes related to primary cilium development (based...
Senescence and Autophagy in Cancer
Enriched terms
0.00
0.04
0.08
0.12
Gene ratio
P-value
0.05
0.10
0.15
0.20
0.25

**S1 Figure. MIC26 loss leads to an opposing regulation of cholesterol biosynthesis pathway upon nutritional stimulation**

(A) Overview of transcriptomics clustering analysis showing upregulated (red) and downregulated (blue) transcripts (without fold change or significance cut-offs). All sample replicates are represented (N = 4).

(B and C) Proteomics data represented by WikPathway enrichment using EnrichR analysis comparing pathways upregulated in normoglycemia (B) and downregulated in hyperglycemia (C) upon *MIC26* deletion compared to respective WT (N = 5). Arrows indicate increased levels of proteins participating in cholesterol synthesis and glycolysis pathways similar to those observed with transcriptomics data (Fig 2C and D). Differentially expressed proteins were considered statistically significant with a cut-off value of fold change of ±1.5 and Bonferroni correction $P \leq 0.05$.
