## Supplementary Figure 2 for "Mitochondrial Apolipoprotein MIC26 is a metabolic rheostat regulating central cellular fuel pathways"

### S2 Figure

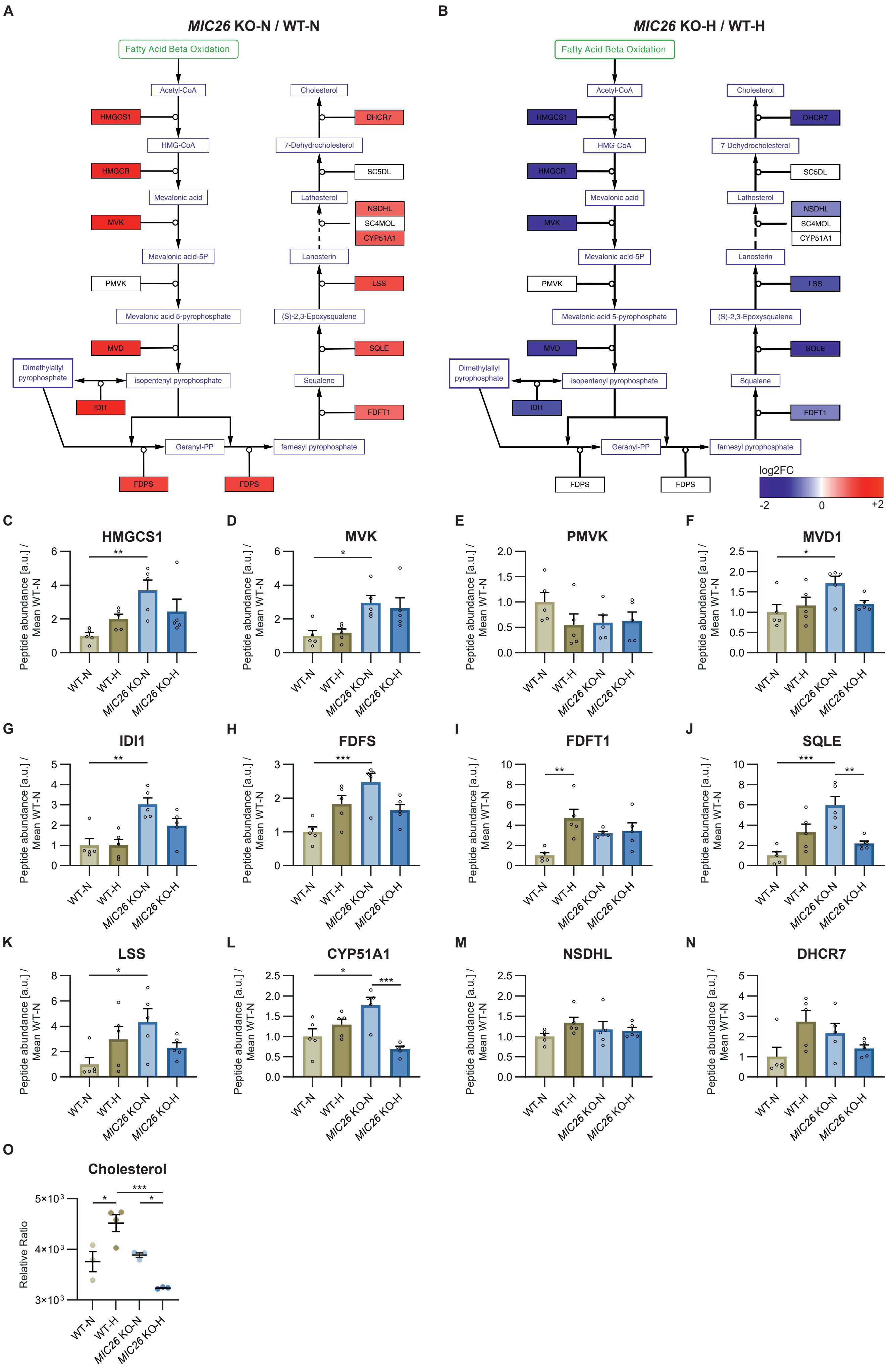
A
MIC26 KO-N / WT-N
B
MIC26 KO-H / WT-H
Fatty Acid Beta Oxidation
Acetyl-CoA
HMGCS1
HMG-CoA
HMGCR
Mevalonic acid
MVK
Mevalonic acid-5P
PMVK
Mevalonic acid 5-pyrophosphate
MVD
Dimethylallyl pyrophosphate
isopentenyl pyrophosphate
IDI1
Geranyl-PP
FDPS
farnesyl pyrophosphate
FDFT1
Squalene
SQLE
(S)-2,3-Epoxysqualene
LSS
Lanosterin
NSDHL
SC4MOL
CYP51A1
Lathosterol
SC5DL
7-Dehydrocholesterol
DHCR7
Cholesterol
log2FC
-2
0
+2
C HMGCS1
D MVK
E PMVK
F MVD1
G IDI1
H FDFS
I FDFT1
J SQLE
K LSS
L CYP51A1
M NSDHL
N DHCR7
Peptide abundance [a.u.] / Mean WT-N
WT-N
WT-H
MIC26 KO-N
MIC26 KO-H
O Cholesterol
Relative Ratio
5×10³
4×10³
3×10³

**S2 Figure. Loss of MIC26 leads to an opposing regulation of cholesterol biosynthesis pathway in normoglycemia and hyperglycemia**

(A and B) The transcripts of various enzymes regulating cholesterol synthesis are represented using Cytoscape software comparing log2FC data of *MIC26* KO and WT cell lines in normo- (A) and hyperglycemia (B) (N = 4). In *MIC26* KO cell lines, normoglycemia strongly increases transcripts of enzymes participating in cholesterol biosynthesis while an opposing effect is observed in hyperglycemia.

(C – N) Peptide abundances of various enzymes participating in cholesterol biosynthesis curated from proteomics data (N = 5).

(O) Metabolomics data reveals that cholesterol levels are exclusively decreased in *MIC26* KO at steady state in hyperglycemia compared to WT but not in normoglycemia (N = 3-4).

Data are represented as mean ± SEM (C-O). Statistical analysis was performed using one-way ANOVA with *$P < 0.05$, **$P < 0.01$, ***$P < 0.001$, ****$P < 0.0001$. N represents the number of biological replicates.
