## Supplementary Figure 3 for "Mitochondrial Apolipoprotein MIC26 is a metabolic rheostat regulating central cellular fuel pathways"

S3 Figure

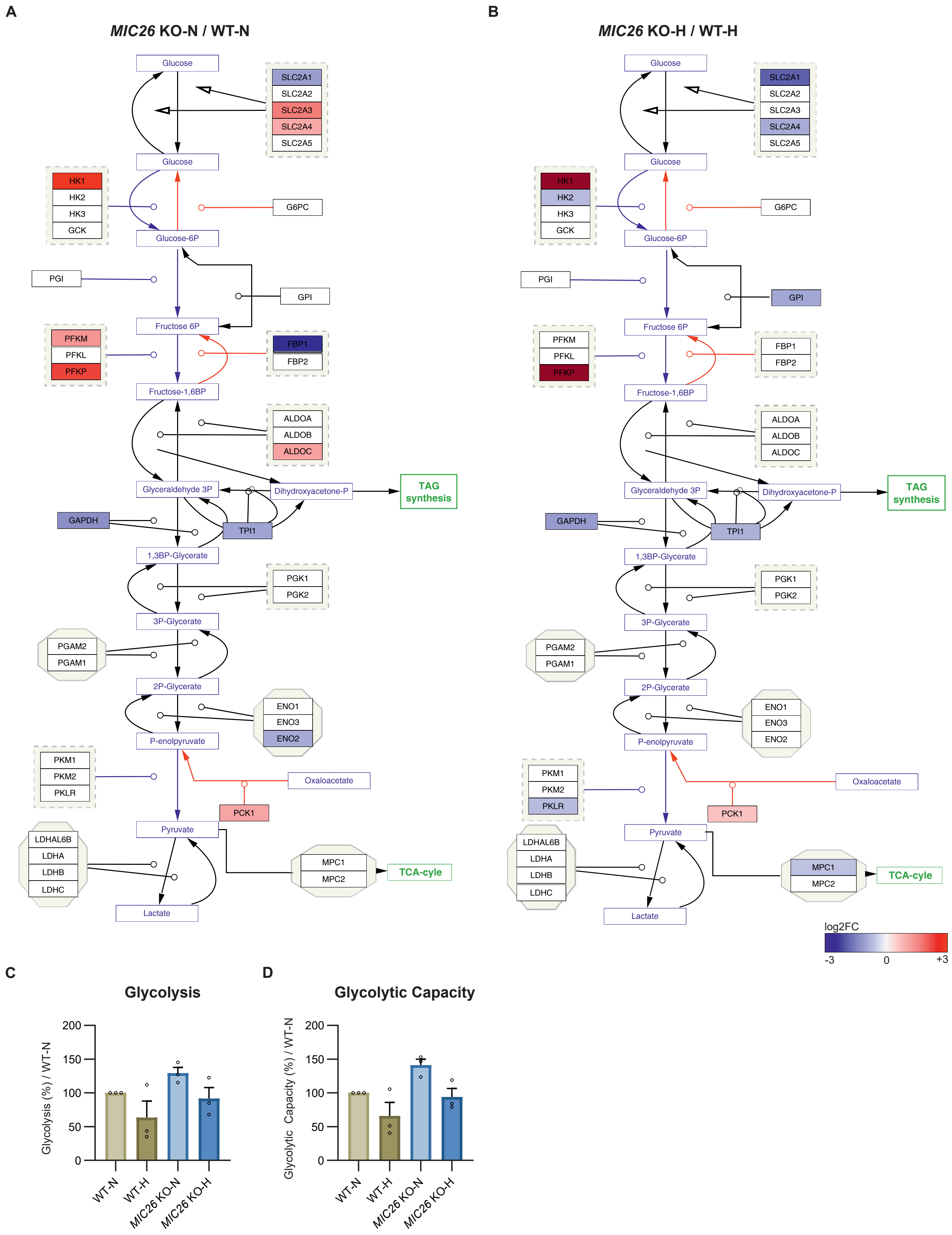
A
MIC26 KO-N / WT-N
B
MIC26 KO-H / WT-H
Glucose
SLC2A1
SLC2A2
SLC2A3
SLC2A4
SLC2A5
HK1
HK2
HK3
GCK
G6PC
Glucose-6P
PGI
GPI
Fructose 6P
PFKM
PFKL
PFKP
FBP1
FBP2
Fructose-1,6BP
ALDOA
ALDOB
ALDOC
Glyceraldehyde 3P
Dihydroxyacetone-P
TAG synthesis
GAPDH
TPI1
1,3BP-Glycerate
PGK1
PGK2
3P-Glycerate
PGAM2
PGAM1
2P-Glycerate
ENO1
ENO3
ENO2
P-enolpyruvate
PKM1
PKM2
PKLR
Oxaloacetate
PCK1
Pyruvate
LDHAL6B
LDHA
LDHB
LDHC
MPC1
MPC2
TCA-cyle
Lactate
log2FC
-3
0
+3
C
Glycolysis
Glycolysis (%) / WT-N
200
150
100
50
0
WT-N
WT-H
MIC26 KO-N
MIC26 KO-H
D
Glycolytic Capacity
Glycolytic Capacity (%) / WT-N

**S3 Figure. *MIC26* deletion causes opposing transcriptional regulation of genes involved in glycolysis**

(A and B) The transcripts of various enzymes participating in glycolysis are represented using Cytoscape software comparing log2FC data of *MIC26* KO and WT cell lines in normo- (A) and hyperglycemia (B) (N = 4).

(C and D) Glycolysis stress test Seahorse assay reveals a nonsignificant tendency towards increased glycolysis and glycolytic capacity upon *MIC26* KO in normoglycemic, but not in hyperglycemic conditions (N = 3).

Data are represented as mean ± SEM (C and D). Statistical analysis was performed using one-way ANOVA with $^{*}P < 0.05$, $^{**}P < 0.01$, $^{***}P < 0.001$, $^{****}P < 0.0001$. N represents the number of biological replicates.
