## Supplementary Figure 4 for "Mitochondrial Apolipoprotein MIC26 is a metabolic rheostat regulating central cellular fuel pathways"

**S4 Figure**

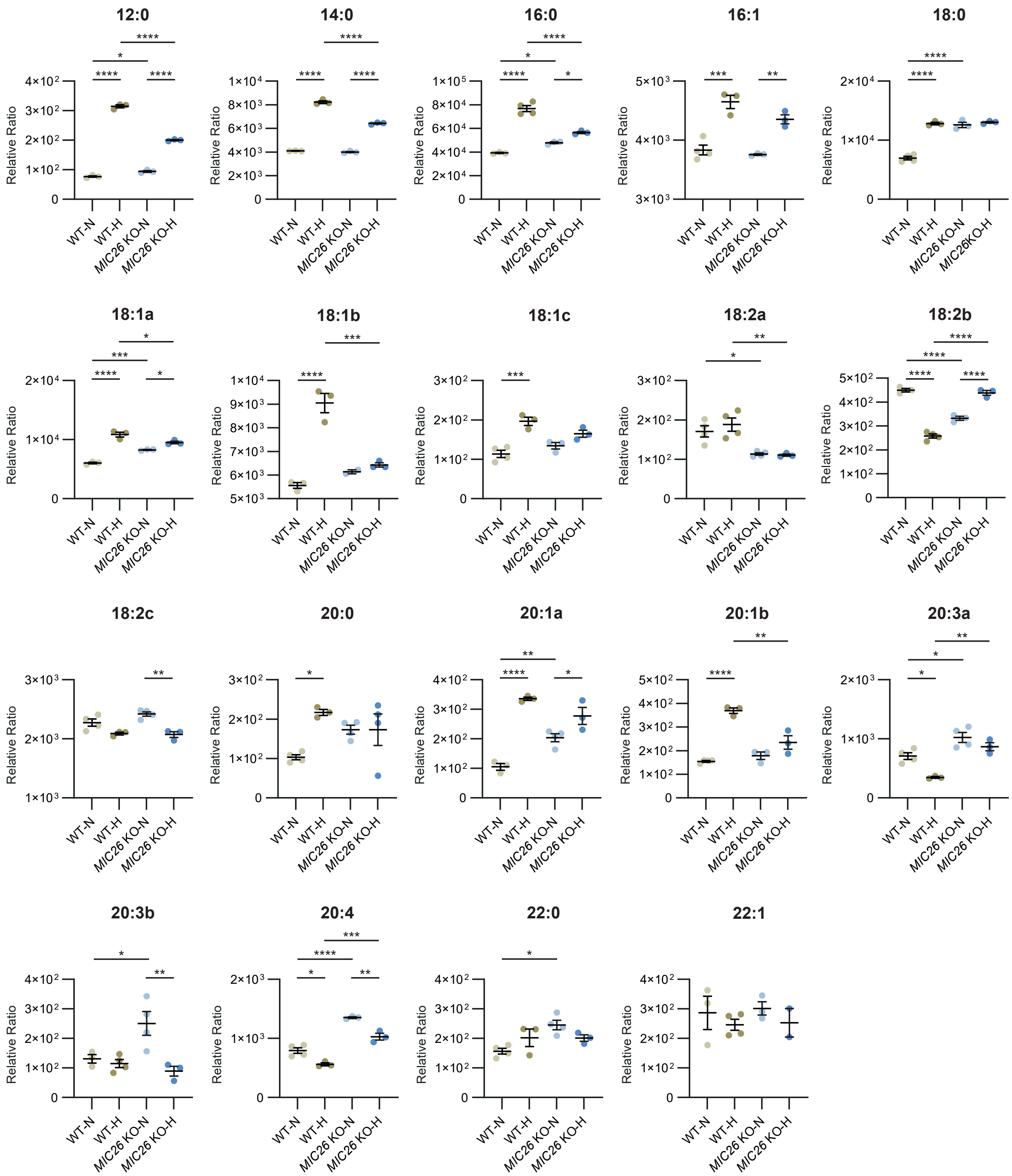

12:0
14:0
16:0
16:1
18:0
18:1a
18:1b
18:1c
18:2a
18:2b
18:2c
20:0
20:1a
20:1b
20:3a
20:3b
20:4
22:0
22:1
Relative Ratio
WT-N
WT-H
MIC26 KO-N
MIC26 KO-H
MIC26KO-H

**S4 Figure. Majority of free fatty acid species are antagonistically regulated upon *MIC26* deletion in normoglycemia and hyperglycemia when compared to respective to WT cells**

Detailed representation of abundances of various free fatty acid species in WT and *MIC26* KO cell lines cultured in normo- and hyperglycemia (N = 3-4).

Data are represented as mean ± SEM. Statistical analysis was performed using one-way ANOVA with *$P < 0.05$, **$P < 0.01$, ***$P < 0.001$, ****$P < 0.0001$. N represents the number of biological replicates.
