## Supplementary Figure 5 for "Mitochondrial Apolipoprotein MIC26 is a metabolic rheostat regulating central cellular fuel pathways"

**S5 Figure**

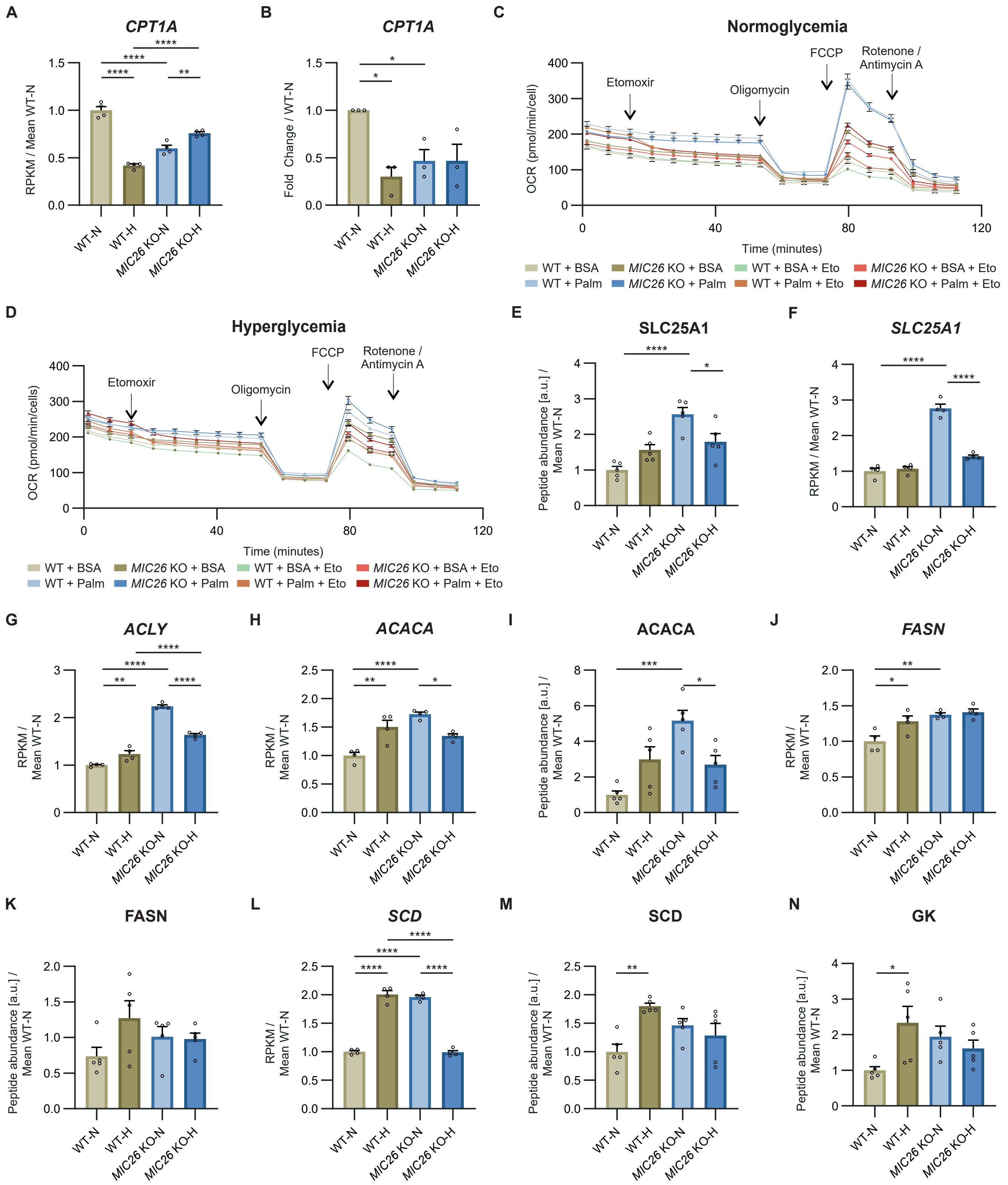

A
CPT1A
RPKM / Mean WT-N
WT-N
WT-H
MIC26 KO-N
MIC26 KO-H
B
CPT1A
Fold Change / WT-N
C
Normoglycemia
Etomoxir
Oligomycin
FCCP
Rotenone / Antimycin A
OCR (pmol/min/cell)
Time (minutes)
WT + BSA
MIC26 KO + BSA
WT + BSA + Eto
MIC26 KO + BSA + Eto
WT + Palm
MIC26 KO + Palm
WT + Palm + Eto
MIC26 KO + Palm + Eto
D
Hyperglycemia
OCR (pmol/min/cells)
E
SLC25A1
Peptide abundance [a.u.] / Mean WT-N
F
SLC25A1
RPKM / Mean WT-N
G
ACLY
RPKM / Mean WT-N
H
ACACA
RPKM / Mean WT-N
I
ACACA
Peptide abundance [a.u.] / Mean WT-N
J
FASN
RPKM / Mean WT-N
K
FASN
Peptide abundance [a.u.] / Mean WT-N
L
SCD
RPKM / Mean WT-N
M
SCD
Peptide abundance [a.u.] / Mean WT-N
N
GK
Peptide abundance [a.u.] / Mean WT-N

**S5 Figure. *MIC26* deletion leads to alteration of key enzymes regulating lipid metabolism**

(A and B) The transcripts of mitochondrial long-chain fatty acid importer CPT1A are strongly decreased in WT-H and *MIC26* KO conditions, compared to WT-N, as shown from transcriptomics data (A) (N = 4) and quantitative PCR (B) (N = 3) analysis.

(C and D) Representative fatty acid oxidation assay analyzed using oxygen consumption rates of WT and *MIC26* KO HepG2 cells cultured in normoglycemia (C) and hyperglycemia (D) upon feeding either with BSA or palmitate (n = 8-12).

(E and F) Mitochondrial citrate malate exchanger (SLC25A1) is significantly increased upon *MIC26* deletion in normoglycemia compared to WT as detected using proteomics (E) (N = 5) and transcriptomics (F) (N = 4) data.

(G – M) Transcripts and available peptide abundances of key genes involved in lipid metabolism curated from transcriptomics (N = 4) and proteomics data (N = 5). Under normoglycemic conditions, loss of MIC26 increases the expression of ATP citrate lyase (G), acetyl-CoA carboxylase 1 (H and I), fatty acid synthase (J and K) and acetyl-CoA desaturase (L and M).

(N) Peptide abundances of glycerol kinase is increased in WT-H compared to WT-N but similar in *MIC26* KO-N and *MIC26* KO-H (N = 5).

Data are represented as mean ± SEM (A-N). Statistical analysis was performed using one-way ANOVA with *$P < 0.05$, **$P < 0.01$, ***$P < 0.001$, ****$P < 0.0001$. N represents the number of biological replicates and n the number of technical replicates.
