## Supplementary Figure 6 for "Mitochondrial Apolipoprotein MIC26 is a metabolic rheostat regulating central cellular fuel pathways"

S6 Figure

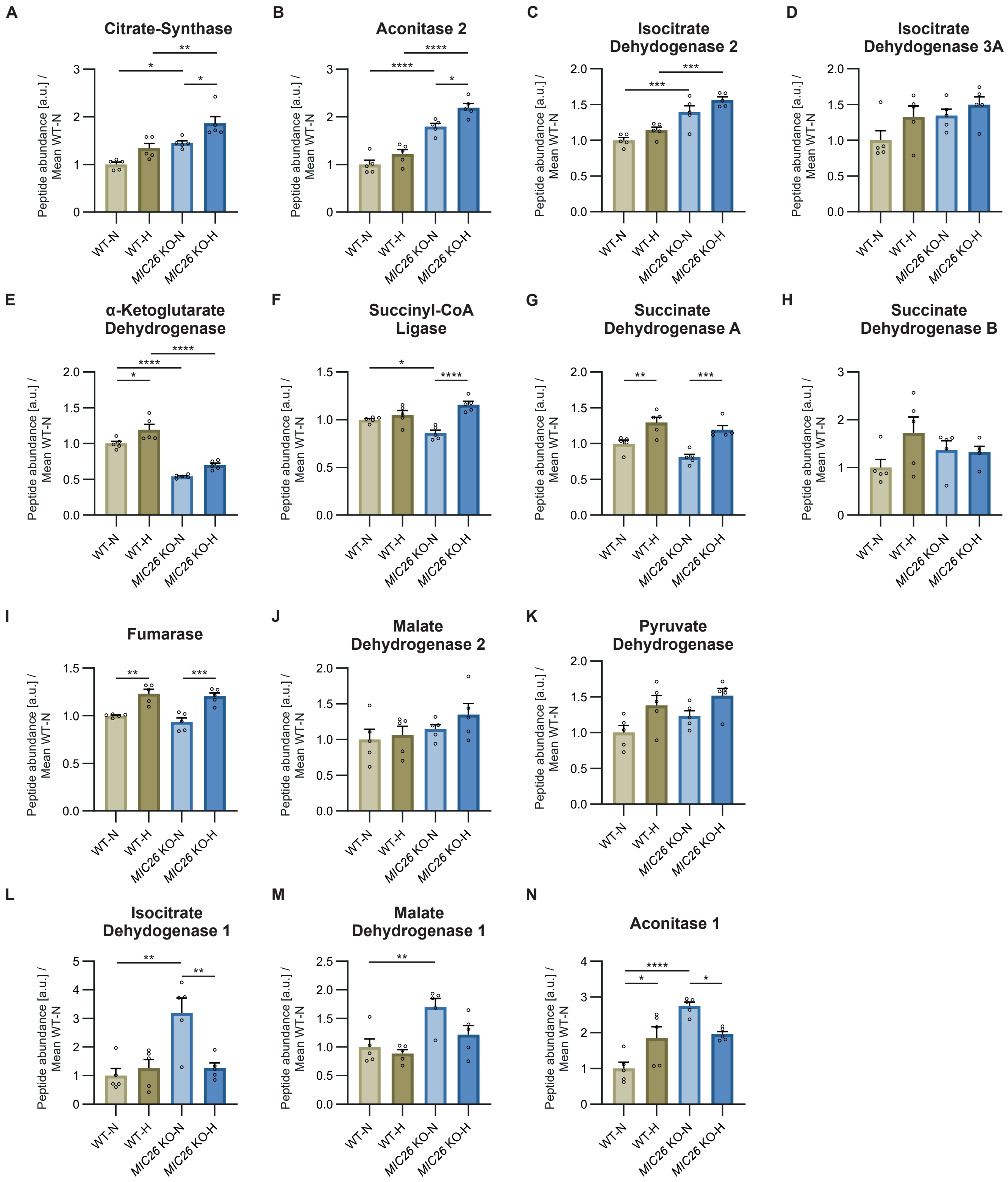
A
Citrate-Synthase
B
Aconitase 2
C
Isocitrate Dehydogenase 2
D
Isocitrate Dehydogenase 3A
E
α-Ketoglutarate Dehydrogenase
F
Succinyl-CoA Ligase
G
Succinate Dehydrogenase A
H
Succinate Dehydrogenase B
I
Fumarase
J
Malate Dehydrogenase 2
K
Pyruvate Dehydrogenase
L
Isocitrate Dehydogenase 1
M
Malate Dehydrogenase 1
N
Aconitase 1
Peptide abundance [a.u.] / Mean WT-N
WT-N
WT-H
MIC26 KO-N
MIC26 KO-H

**S6 Figure. TCA cycle enzymes are altered upon *MIC26* knockout**

(A – K) Representation of peptide abundances of various mitochondrial TCA cycle enzymes curated from proteomics data (N = 5).

(L – N) Peptide abundances of cytosolic enzymes involved in metabolite conversion (N = 5).

Data are represented as mean ± SEM (A-N). Statistical analysis was performed using one-way ANOVA with $^*P < 0.05$, $^{**}P < 0.01$, $^{***}P < 0.001$, $^{****}P < 0.0001$. N represents the number of biological replicates.
