## Supplementary Figure 7 for "Mitochondrial Apolipoprotein MIC26 is a metabolic rheostat regulating central cellular fuel pathways"

### S7 Figure

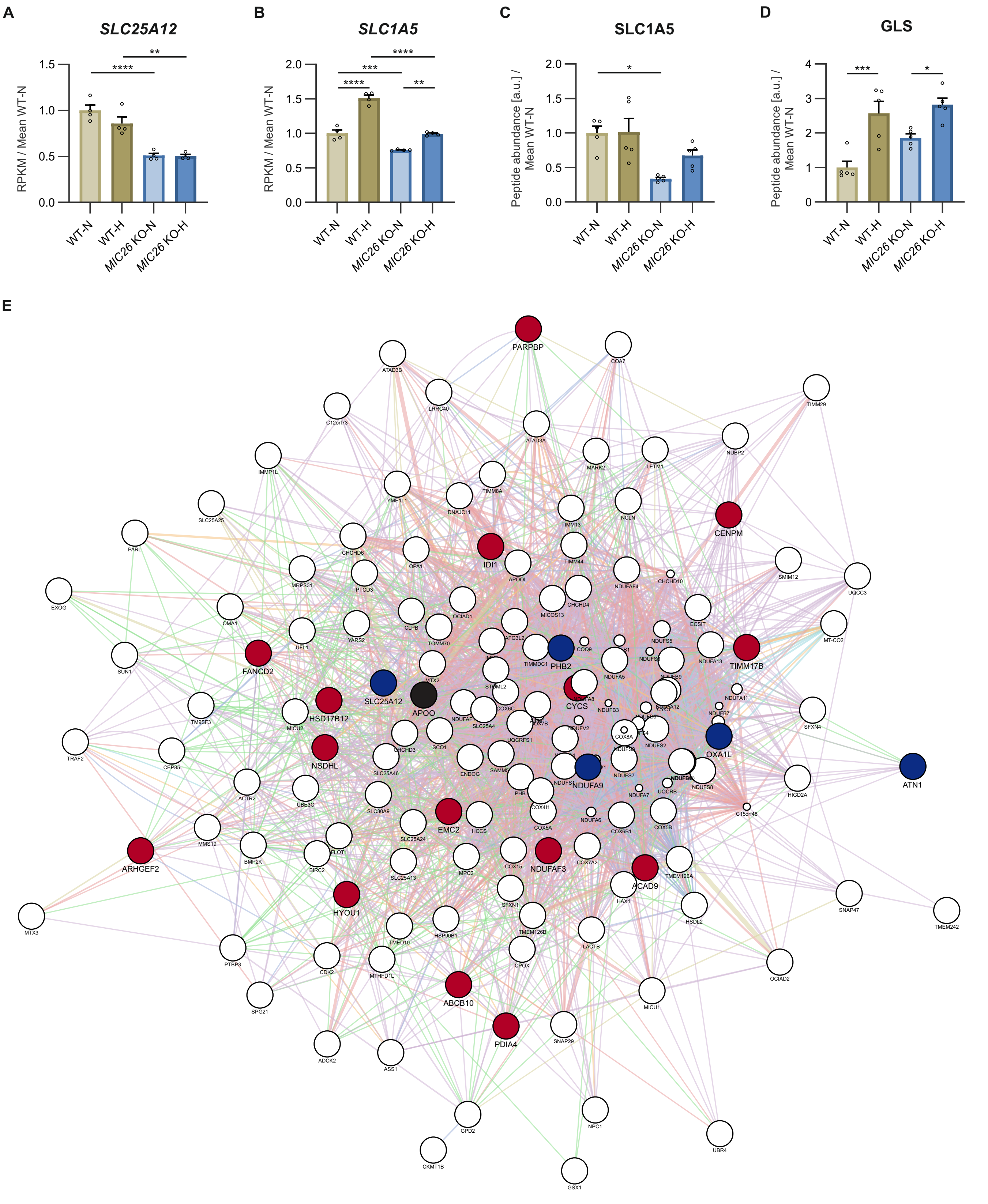
A
SLC25A12
RPKM / Mean WT-N
1.5
1.0
0.5
0.0
****
**
WT-N
WT-H
MIC26 KO-N
MIC26 KO-H
B
SLC1A5
RPKM / Mean WT-N
2.0
1.5
1.0
0.5
0.0
****
***
****
**
WT-N
WT-H
MIC26 KO-N
MIC26 KO-H
C
SLC1A5
Peptide abundance [a.u.] / Mean WT-N
2.0
1.5
1.0
0.5
0.0
*
WT-N
WT-H
MIC26 KO-N
MIC26 KO-H
D
GLS
Peptide abundance [a.u.] / Mean WT-N
4
3
2
1
0
***
*
WT-N
WT-H
MIC26 KO-N
MIC26 KO-H
E
PARPBP
CENPM
IDI1
TIMM17B
FANCD2
PHB2
SLC25A12
APOO
CYCS
HSD17B12
OXA1L
ATN1
NSDHL
NDUFA9
EMC2
ARHGEF2
NDUFAF3
ACAD9
HYOU1
ABCB10
PDIA4

**S7 Figure. Mitochondrial glutamine and glutamate carriers are downregulated upon loss of MIC26**

(A) Transcripts of mitochondrial glutamate aspartate antiporter *SLC25A12* (A) are decreased upon *MIC26* deletion in normo- and hyperglycemia (N = 4).

(B and C) Transcripts (B) (N = 4) and peptide abundances (C) (N = 5) of cellular and mitochondrial glutamine importer SLC1A5 are significantly decreased upon *MIC26* deletion in both normo- and hyperglycemia in relation to respective WT cells.

(D) Peptide abundances of glutaminase (GLS) are unaltered upon loss of MIC26 compared to respective WT conditions.

(E) MIC26 interactome, based cumulatively on BioGRID, NeXtProt and IntAct databases, generated with Cytoscape software. From this study, downregulated transcripts comparing *MIC26* KO and WT in normoglycemic condition are highlighted in blue while upregulated transcripts are highlighted in red.

Data are represented as mean ± SEM (A-D). Statistical analysis was performed using one-way ANOVA with *$P < 0.05$, **$P < 0.01$, ***$P < 0.001$, ****$P < 0.0001$. N represents the number of biological replicates.
