## Supplementary Figure 8 for "Mitochondrial Apolipoprotein MIC26 is a metabolic rheostat regulating central cellular fuel pathways"

S8 Figure

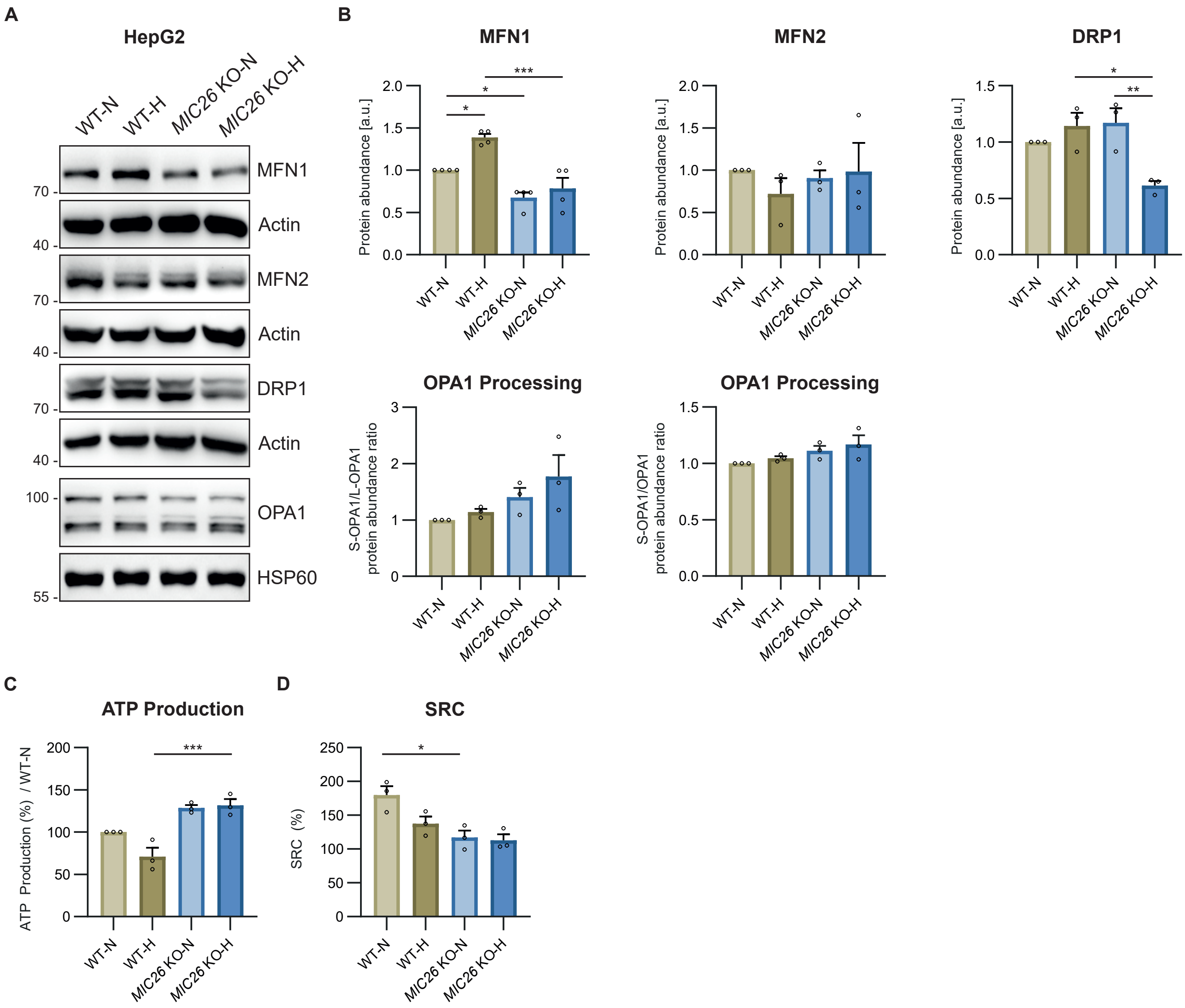
A
HepG2
WT-N
WT-H
MIC26 KO-N
MIC26 KO-H
MFN1
Actin
MFN2
Actin
DRP1
Actin
OPA1
HSP60
70
40
70
40
70
40
100
55
B
MFN1
MFN2
DRP1
Protein abundance [a.u.]
OPA1 Processing
S-OPA1/L-OPA1 protein abundance ratio
S-OPA1/OPA1 protein abundance ratio
C
ATP Production
ATP Production (%) / WT-N
D
SRC
SRC (%)

**S8 Figure. MIC26 maintains mitochondrial morphology and bioenergetics**

(A and B) Western Blots (A) and quantification (B) show a decrease of key mitochondrial fusion mediator MFN1 in *MIC26* KO cells while MFN2 was unchanged. Mitochondrial fission mediator DRP1 is decreased in *MIC26* KO-H compared to WT-H. OPA1 processing shows no significant changes upon *MIC26* deletion and nutritional status (N = 3-4).

(C and D) ATP production (C) and spare respiratory capacity (D) determined by mito stress test using Seahorse XF analyzer. Deletion of *MIC26* caused increased ATP production and decreased metabolic flexibility (indicated by SRC) in normoglycemia (N = 3).

Data are represented as mean ± SEM (B-D). Statistical analysis was performed using one-way ANOVA with *$P < 0.05$, **$P < 0.01$, ***$P < 0.001$, ****$P < 0.0001$. N represents the number of biological replicates.
