## Supplementary Figure 9 for "Mitochondrial Apolipoprotein MIC26 is a metabolic rheostat regulating central cellular fuel pathways"

S9 Figure

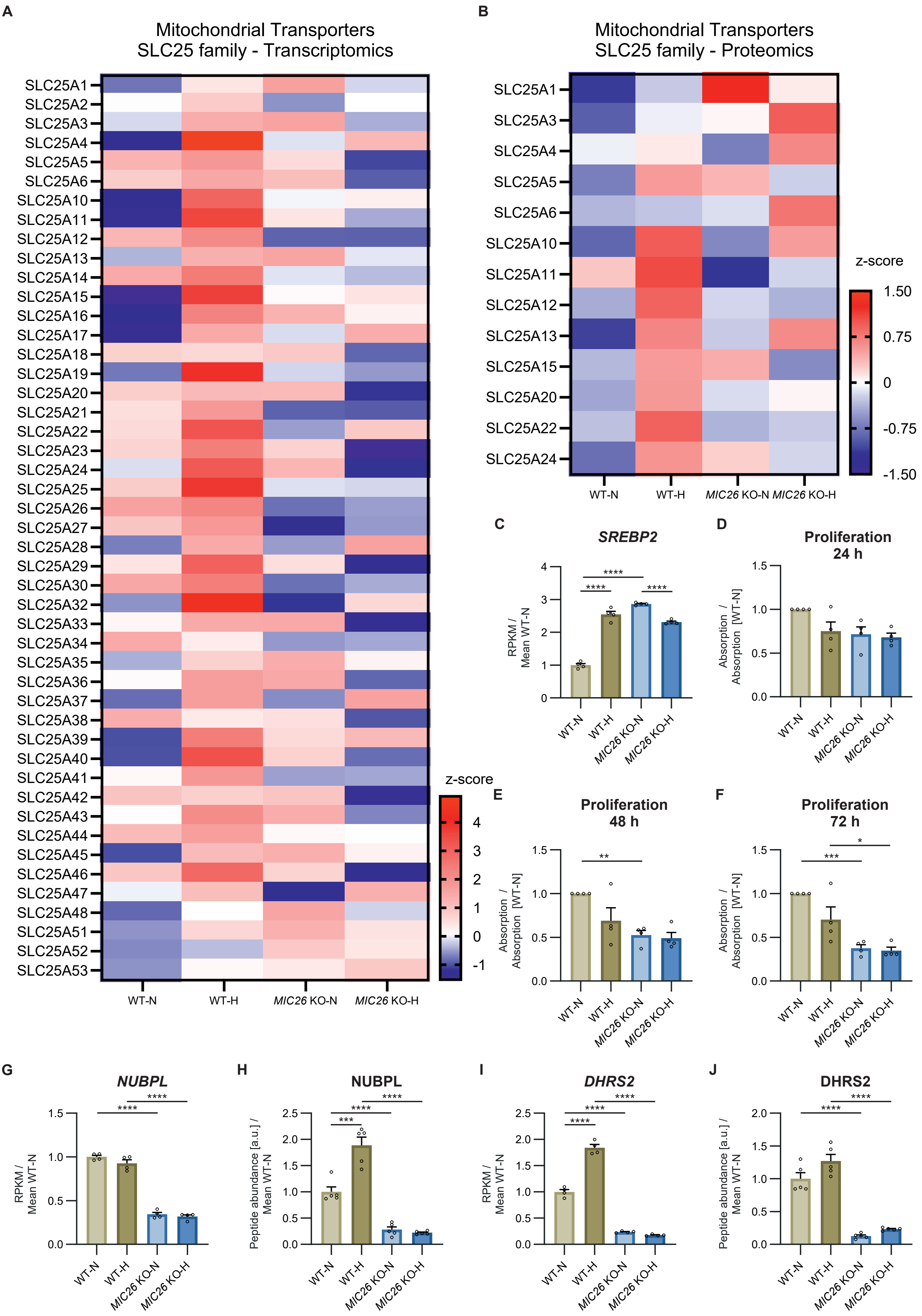
A
Mitochondrial Transporters
SLC25 family - Transcriptomics
SLC25A1
SLC25A2
SLC25A3
SLC25A4
SLC25A5
SLC25A6
SLC25A10
SLC25A11
SLC25A12
SLC25A13
SLC25A14
SLC25A15
SLC25A16
SLC25A17
SLC25A18
SLC25A19
SLC25A20
SLC25A21
SLC25A22
SLC25A23
SLC25A24
SLC25A25
SLC25A26
SLC25A27
SLC25A28
SLC25A29
SLC25A30
SLC25A32
SLC25A33
SLC25A34
SLC25A35
SLC25A36
SLC25A37
SLC25A38
SLC25A39
SLC25A40
SLC25A41
SLC25A42
SLC25A43
SLC25A44
SLC25A45
SLC25A46
SLC25A47
SLC25A48
SLC25A51
SLC25A52
SLC25A53
WT-N
WT-H
MIC26 KO-N
MIC26 KO-H
z-score
4
3
2
1
0
-1
B
Mitochondrial Transporters
SLC25 family - Proteomics
SLC25A1
SLC25A3
SLC25A4
SLC25A5
SLC25A6
SLC25A10
SLC25A11
SLC25A12
SLC25A13
SLC25A15
SLC25A20
SLC25A22
SLC25A24
WT-N
WT-H
MIC26 KO-N
MIC26 KO-H
z-score
1.50
0.75
0
-0.75
-1.50
C
SREBP2
RPKM / Mean WT-N
****
****
****
D
Proliferation 24 h
Absorption / Absorption [WT-N]
E
Proliferation 48 h
Absorption / Absorption [WT-N]
**
F
Proliferation 72 h
Absorption / Absorption [WT-N]
***
*
G
NUBPL
RPKM / Mean WT-N
****
****
H
NUBPL
Peptide abundance [a.u.] / Mean WT-N
***
****
****
I
DHRS2
RPKM / Mean WT-N
****
****
****
J
DHRS2
Peptide abundance [a.u.] / Mean WT-N
****
****

**S9 Figure. *MIC26* deletion induces alterations of SLC25 mitochondrial carrier protein family expression and induces growth defects**

(A and B) Heat map overview from mean z-score of transcripts (A) (N = 4) and peptide abundances (B) (N = 5) of mitochondrial transporters belonging to SLC25 family.

(C) Transcript abundances of *SREBP2* (C) of WT and *MIC26* KO cells grown in normo- and hyperglycemia (N = 4).

(D – F) Proliferation of respective cell lines after 24 h (C), 48 h (D) and 72 h (E) determined using SRB assay normalized to WT-N (N = 4).

(G – J) Transcripts (N = 4) and peptide abundances (N = 5) of NUBPL (G and H) and DHRS2 (I and J) are respectively shown.

Data are represented as mean ± SEM (C-J). Statistical analysis was performed using one-way ANOVA with *$P < 0.05$, **$P < 0.01$, ***$P < 0.001$, ****$P < 0.0001$. N represents the number of biological replicates.
